# Comparative single-cell transcriptional atlases of *Babesia* species reveal conserved and species-specific expression profiles

**DOI:** 10.1101/2022.02.11.480160

**Authors:** Yasaman Rezvani, Caroline D Keroack, Brendan Elsworth, Argenis Arriojas, Marc-Jan Gubbels, Manoj T Duraisingh, Kourosh Zarringhalam

## Abstract

*Babesia* is a genus of Apicomplexan parasites that infect red blood cells in vertebrate hosts. Pathology occurs during rapid replication cycles in the asexual blood-stage of infection. Current knowledge of *Babesia* replication cycle progression and regulation is limited and relies mostly on comparative studies with related parasites. Due to limitations in synchronizing *Babesia* parasites, fine-scale time-course transcriptomic resources are not readily available. Single-cell transcriptomics provides a powerful unbiased alternative for profiling asynchronous cell populations. Here, we applied single-cell RNA sequencing to three *Babesia* species (*B. divergens, B. bovis*, and *B. bigemina)*. We used analytical approaches and algorithms to map the replication cycle and construct pseudo-synchronized time-course gene expression profiles. We identify clusters of co-expressed genes showing *just-in-time* expression profiles, with gradually cascading peaks throughout asexual development. Moreover, clustering analysis of reconstructed gene curves reveals coordinated timing of peak expression in epigenetic markers and transcription factors. Using a regularized Gaussian Graphical Model, we reconstructed co-expression networks and identified conserved and species-specific nodes. Motif analysis of a co-expression interactome of AP2 transcription factors identified specific motifs previously reported to play a role in DNA replication in *Plasmodium* species. Finally, we present an interactive web-application to visualize and interactively explore the datasets.

## INTRODUCTION

Apicomplexan parasites of the genus *Babesia* are some of the most widespread blood parasites of vertebrates, second only to the trypanosomes (1). *Babesia* has long been recognized as a disease of tremendous veterinary and agriculture importance, causing hundreds of millions of dollars of economic losses every year (2, 3). Since the first reported case of human babesiosis, caused by *B. divergens* reported in 1956, compounded by the emergence of *B. microti* in the USA, babesiosis has steadily been gaining recognition as an important human parasitic disease (4–7). The disease can range from mild febrile illness to severe, life-threatening disease, particularly in immunocompromised patients (6, 8). *Babesia* is usually transmitted through the bite of an infected tick (9), but can also be transmitted congenitally and via blood transfusion (10–12). Indeed, *Babesia* is listed as a top priority pathogen in the blood supply (13). Beyond human pathogens, there are at least 100 species of *Babesia* described which cause disease in a variety of hosts (6, 14). Indeed, bovine babesiosis, predominantly caused by *B. bovis*, *B. bigemina*, and *B. divergens*, is of significant concern, often resulting in fulminating infection and high mortality, leading to significant economic and agricultural losses (15, 16). With such a wide diversity of disease-causing parasites, identifying both conserved and divergent biology is essential to developing therapeutic and vaccine interventions.

In the asexual replicative cycle, *Babesia* are obligate intracellular parasites that infect red blood cells (RBCs). While knowledge about the morphology of *Babesia* parasites during these division cycles exists, molecular details of the asexual replication cycle is limited. Transcriptomic studies have been done on various *Babesia* spp. populations to profile life stage, egress and invasion, and virulence (17–22). While these studies provide rich data resources, no transcriptomic data have been generated to comprehensively describe the asexual replication cycle, as has been done in other related parasites (23–27). Most knowledge of the molecular mediators of the *Babesia* spp. replication cycle has been gleaned through comparative approaches with *Plasmodium* spp. and *T. gondii* (28), and through the analysis of genomic sequences (17, 18, 29–36). Likely due to the difficulty in obtaining synchronous populations of parasites during intraerythrocytic development, to date only a single synchronous transcriptomic dataset exists (37). Additionally, no single-cell RNA sequencing (scRNA-seq) has yet been performed in any *Babesia* species.

Due to these gaps in knowledge, and the limitations of synchronization, scRNA-seq presents as a promising method for delineating the *Babesia* intraerythrocytic replication cycle at a fine scale. This approach offers a powerful, unbiased method of profiling heterogenous and asynchronous cell populations. Single-cell RNA-seq has been used successfully in a range of other apicomplexan parasites and has provided key insights into parasite biology not previously detectable using bulk RNA sequencing methods. In *Plasmodium* species, scRNA-seq has been used to describe cell populations through the entire life cycle, from the IDC through the mosquito (38–45). This has led to the development of comprehensive cell atlases for both *P. berghei* (39) and *P. falciparum* (46). Additionally, scRNA-seq in *T. gondii* was used to reveal novel regulators of life cycle stage progression (47, 48). Combined with the ability to culture many diverse species of *Babesia in vitro* (49, 50), comparative scRNA-seq offers a unique opportunity in *Babesia* to identify core, conserved regulators of progression through the replicative cycle.

Here, we performed scRNA-seq on three bovine *Babesia* species that can be readily cultured *in vitro*: *B. bovis, B. bigemina*, and *B. divergens.* Additionally, to investigate a possible role of the host RBC on the replication cycle, we performed scRNA-seq on *B. divergens* adapted to either human RBCs or bovine RBCs in *in vitro* culture. Using these four data sets, we reconstructed a transcriptome of the asexual replication cycle for *Babesia* spp. and mapped the transition points of the developmental phases. We show that despite their evolutionary divergence, the underlying expression pattern of the replication cycle in *Babesia* is highly similar. Using these data, we were also able to reconstruct gene co-expression networks and quantify interactomes of gene families important in each phase of development. Analysis of the interactome reveals a core conserved set of replication cycle regulators. Taken together, these data represent the first asexual replication cycle atlases for any *Babesia* species, as well as the first single-cell transcriptomic data sets. To facilitate usage, we provide an interactive web-application for visualization and exploration of our datasets. The web-app provides functionality to profile gene-expression, performed comparative transcriptomics analysis, assess and visualize gene-expression timing across the species, and explore co-expression networks. The web-app is hosted at: https://umbibio.math.umb.edu/babesiasc/.

## MATERIALS AND METHODS

### Parasite culture

The *Babesia bovis* strain MO7 and the *B. bigemina* strain J29, provided by David Allred of the University of Florida, were maintained in purified bovine RBCs (hemostat) hematocrit in RPMI-1640 media supplemented with 25 mM HEPES, 11.50 mg/l hypoxanthine, 2.42 mM sodium bicarbonate, and 4.31 mg/ml AlbuMAX II (Invitrogen). Before addition of AlbuMAX-II and sodium bicarbonate, we adjusted the pH of the media to 6.75. *Babesia divergens* strain Rouen 1987, kindly provided by Kirk Deitsch and Laura Kirkman (Weill Cornell Medical College), maintained under the same conditions in purified Caucasian male O+ human RBCs (Research Blood components). All cultures were maintained at 37 °C in a hypoxic environment (1% O_2_, 5% CO_2_). Clonal lines of parasites were used for all selections and were derived from the provided strains via limiting dilution-these will be referenced as BdC9 (*B. divergens*), BigE1 (*B. bigemina*), and BOV2C (*B. bovis*).

### Single-cell protocol

*B. divergens, B. bovis*, and *B. bigemina* were grown to >15% parasitemia. Health of parasites was assessed by thin blood smear. Parasites were collected and pelleted, and washed with warm 1X PBS, followed by a final wash with warm 0.4% BSA in 1X PBS. To ensure loading of the correct number of parasites, parasitemia was counted on stained thin blood smears (2000 total cells), and red blood cells were counted using a hemocytometer. These values were used to calculate the number of infected cells to be loaded into the Chromium Chip B. We aimed for recovery of 10, 000 infected cells, thus loaded 16, 500 infected cells in bulk culture. Cell suspensions were loaded into individual wells on the Chromium Chip B. Post gel beads-in-emulsion (GEM) generation, single-cell libraries were processed according to the 10 × Chromium 3′ v2 User Guide protocol, using 13 amplification cycles for cDNA amplification, and 14 cycles in library construction. Libraries were subsequently sequenced on the Illumina NextSeq platform following the 10X specifications, aiming for a minimum of 6000 reads per cell.

### RNA-seq alignment

The reference genomes and annotation files of *Babesia* spp. (release 54) were downloaded from https://piroplasmadb.org/. Custom references were generated using the 10X cellranger pipeline (cellranger mkref) and raw fastq files were aligned to the genome cellranger count with default parameters.

### Single-Cell RNA-seq data processing

The R Seurat package (51) was utilized to process the count data. Seurat objects were created for each count data independently. Cells and genes with low counts were filtered out from the analysis using the Seurat function CreateSeuratObject with parameters min.cells = 10, and min.features = 100. Expression data was normalized using Seurat functions FindVariableFeatures with parameter nfeatures = 3000 and ScaleData. Dimensionality reduction was performed using principal component analysis (PCA) and Uniform Manifold Approximation & Projection (UMAP) as implemented in Seurat functions RunPCA, and FindNeighbors, with parameters dims = 1:10 and reduction = ‘pca’. Clustering analysis was performed using k-nn algorithm using Seurat function FindClusters with parameter res = 0.2. Datasets were down sampled to include 800 cells per cluster. Orthologous genes in all three species were used to construct Seurat objects with same genes using *B. divergens* gene ids. Datasets were then integrated using Seurat’s merge and IntegrateData functions. The *B. divergens* sample in human host RBCs was used as the ‘reference’ dataset for integration (Seurat FindIntegrationAnchors function).

### Pseudo-time analysis

Pseudo-time analysis was performed in 3 steps. First, an ellipsoid was fitted to the first two PCA coordinates in each dataset using Ellipsefit function from MyEllipsefit R package (https://github.com/MarkusLoew/MyEllipsefit). Next, the elliptic fit was used as prior to fit a principal curve to each data set (52).The function principal_curve from the R package princurve was used to fit the principal curves. The parameter smoother = “periodic_lowess” was set to enforce closed curves. Data was then orthogonally projected onto the principal curves and ordered to generate pseudo time. The pseudo time curve was mapped to interval [0, 12 h] to mimic the *Babesia spp.* replication cycle. The pseudo-time interval was partitioned into 20 min bins and cells that fell within the same bin were treated as synchronized replicates. Next, we calculated the correlation of gene expression with pseudo-time using a Generalized Additive Model (GAM) and filtered out genes that did not correlate with the pseudo-time (FDR adjusted p-value < 0.05). The R function gam from the package gam was used for this analysis. This resulted in a time-course expression matrix with dimensions *n × ∏^N^ _k = 1_ n_k_* with *n* representing the total number of genes, *N* representing total number of time bins (36 intervals, each 20 min), and *n_k_* representing total number of cells mapping to the time bin *k*.

### Fitting gene curves: Mixed-effect model

The following mixed-effect smoothing spline model was used to estimate mean-expression of genes along the pseudo-time:

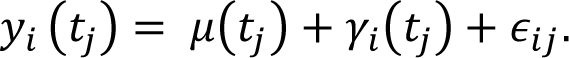

Here *y_i_(t_j_)* represents the expression of gene *i* in the binned time interval *j, μ(t)* is the fix effect corresponding to population mean, *γ_i_(t)* stands for the random effect corresponding to deviation of gene expression from mean at each time point and *∈_ij_* is the assumed independent normally distributed noise. As before cells within the same time partition were considered replicates. The random effect is from a Gaussian Process γ_i_ ∼ GP(0, δ) with mean 0 and covariance matrix *D(ℓ, S) = δ(t_ℓ_, t_S_)*. The function sme() function from the R package sme was used to fit this model. Next, a natural cubic spline was fitted to the coefficients returned by sme() and gene curves were interpolated at regular time intervals. The fitted objects were used to visualize the mean curve along with the confidence bands at 95% level.

### Alignment with bulk RNA-seq data

Time course Bulk RNA-seq data from synchronized *B. divergens* parasite was processes as previously described (37). The bulk time course data consisted of 7 measurements of synchronized *B. divergens* parasites over 12 h period spanning the replication cycle. There was a total of 2 biological replicates and 2 technical replicates. Technical replicates were merged for this analysis. Smoothing splines were fitted to the common genes between the bulk time-course data and single-cell pseudo-time course data. The smoothing spline fits to the bulk data were samples to 36 points, matching the total number of points in the single-cell gene curve data. Cross correlation between corresponding genes in bulk and *B. divergens* single-cell datasets was calculated by

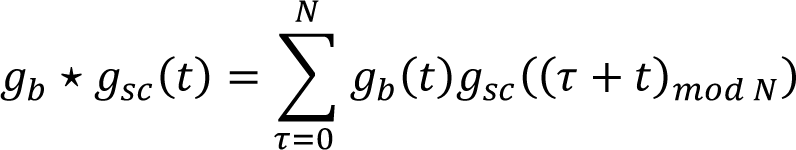

where *g_b_(t) and g_SC_ (t)* are gene curves is the bulk and singe cell data respectively and *N* is the total number of sample points in the gene curves. Lag-time maximizing the cross-correlation was then calculated for each gene and the distribution of lag-times across all genes was examined to identify a single optimal lag-time for all genes. Single-cell gene curves in all single-cell datasets were shifted by the optimal lag time to adjust the start time.

### Inferred Replication cycle Phases

*T. gondii* scRNA-seq data was obtained from the single-cell atlas of *T. gondii* (48), where replication cycle phases were determined using DNA content and computational analysis. Markers of each phase were determined by performing differential expression analysis using Seurat R function findAllMarkers with parameters only.pos = TRUE, min.pct = 0. Significance was determined using fold change > 2 and adjusted p-value < 0.01. Top 20 markers of each phase were then used and mapped to their *Babesia spp.* orthologs. Timing of peak expression of each marker was calculated by examining the local maxima of the fitted pseudo-time gene expression curves in each species. Transition time points between phases were then determined by examining the quantiles of the peak time distributions and adjusted by visual inspection of the overlap of distributions.

### Marker analysis

Markers of the inferred replication cycle phases were identified as follows. Conserved markers of each phase across species were identified using the Seurat function FindConservedMarkers default parameters. Markers of each phase were also calculated in each species independently using FindAllMarkers function from the Seurat R package with parameters only.pos = TRUE. Markers of each phase unique to a specific species were determined using the same function and by setting an appropriate contrast in cell identities (e.g. *B. bigemina* G phase vs *B. bovis*, *B. divergens* (bovine), *B. divergens* (human) G phase). For these analyses fold change cutoff > 2 and adjusted p-value < 0.01 were used to determine significance. For host cell specific differences, differentially expression analysis was performed between *B. divergens* in human host and *B. divergens* in bovine host as well as *B. divergens* in human host and merged *B. bigemina*, *B. bovis*, *B. divergens* in bovine host in phased matched specific manner. For these analyses fold change cutoff > 1.5 and adjusted p-value < 0.01 were used to determine significance.

### Enrichment analyses

Differentially expressed genes were mapped to their orthologs in *T. gondii* orthologs. Gene Ontology Enrichment Analysis (GOEA) was performed using available GO on ToxoDB.org and significant GO terms (Benjamini < 0.1) were determined. The log (fold enrichment) and log (p-value) were used to visualize the significant GO terms.

### Time-course clustering

Mean gene expressions curves of inferred replication cycle marker genes were used to construct an *n × N* time-course matrix, with rows representing genes and columns representing pseudo-time bins. Data was scaled to z-scores and a Dynamic Time-Warping (DTW) metric was used to measure the similarity between curves and perform a hierarchical clustering. For this analysis we used the tsclust function from the R package dtwclust (https://github.com/asardaes/dtwclust) with parameters control = hierarchical_control(method = “complete”), args = tsclust_args(dist = list(window.size = 4L). Total number of clusters were set empirically with trial and error. Genes were ordered according to peak expression time and cells were according to their inferred replication cycle phase (transition points along the pseudo-time). A heatmap was used to visualize the expression profile of the genes.

### Reconstruction of gene-gene interaction network

We used a Gaussian Graphical Model to assemble a gene-gene interaction network using the scRNA-seq expression data. The GMM can be used to calculate the partial correlation between gene-pairs, conditioned on the rest of the genes and as such, it captures pairwise relationships between the nodes in the interaction graph. Partial correlation is then used to assemble a gene-gene interaction network, where genes represent nodes and edges represent a direct interaction between them after accounting for tertiary effects. The objective of the GGM is given by

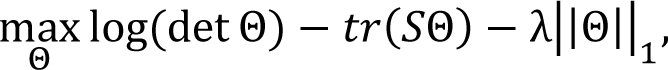

where *S* and Θ are the empirical covariance and precision matrices and λ is the sparsity penalty. To fit this model, we estimated the empirical covariance matrix S using the single pseudo-time course gene expression data as follows. First, the mean trend μ_i_(t_j_) of each gene i at time point t_j_ was estimated using the expression of replicate cells that mapped to time partition i. This mean trend was removed from the expression of genes to de-trend the data:

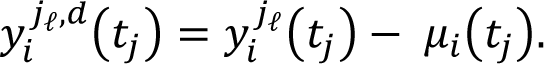

The superscript represents (replicate) cell j_ℓ_ at time bin j. The trended data was used to estimate the empirical covariance matrix S. As gene expression is periodic during the replication cycle, and assuming a non-time varying covariance matrix, the GMM model can be directly applied to the de-trend data to capture direct covariations in gene expression. The Objective function of the GMM was then fitted for a grid of λ values ranging from 0.01 to 1.0 with step-size 0.01. The R package glassoFast was used for fitting the model (https://github.com/JClavel/glassoFast). The fitted precision matrices were converted to partial correlation matrices Ρ, which in turn were converted to network adjacency matrices. The scale free network property for each network was calculated and the penalty value that maximized the scale free property was used to identify the “optimal” network.

### Motif search analysis

Annotated *P. falciparum* TFs and AP2s were mapped to their *Babesia spp.* orthologs when available (**SI** table 4). For each TF, the interactomes were identified in the assembled co-expression networks in each species and the union of all interacting genes across all species was taken as the TF’s overall interactome. The promoter of genes in the overall interactomes of each TF were defined as the sequence on N nucleotide upstream of the start codon, where N was set to 167 in *B. bigemina*, 219 in *B. bovis*, and 190 in *B. divergens*. These promoter lengths were selected as 1/2 of the mean gap between consecutive genes in each species, calculated from genome annotation files. The promoters were extracted using the getfasta command from bedtools package (53) and the latest versions of the genome of each species, downloaded from https://piroplasmadb.org/. Motif search analysis was carried out on promoter sequences using MEME suite (54). The meme command with the following parameters: -dna -minw 4 -maxw 10 anr -nmotifs 10 -revcomp was used to perform the motif search.

### Data visualization

A web dashboard was built to provide easy access to single-cell expression data and analysis results. Data was preprocessed and loaded as tables to a SQL database. The web interface was implemented using the ‘dash’ python framework, which allows to build dynamic, interactive, data-driven apps. The current instance of the app is running in a single server using docker containers. However, the app design and the stateless approach of the framework allows for easy scalability to support increasing traffic as needed. The app for the dashboard is organized as a python module and separated in a sub-module for each of the pages available. URL requests are processed using an index python script and loads the appropriate UI layout and backend logic. The python app is served by a guinicorn WSGI HTTP Server, which allows to spawn multiple workers for the app. MariaDB is used as the SQL server for the app, with connection pools of size 32 for each python worker, allowing multiple persistent connections to the database. A series of bash and python scripts were written to automate the process of database initialization from the TSV datafiles. As the data is expected to remain static through the running time of the app, SQL tables were created with tight data size allocations to help with performance. SQL indices were created such that queries remain fast. SQL unique indices are used where possible, as a way of ensuring the input data’s integrity. The resulting relational database contains 9 tables for each of the species, holding data and meta-data for genes, orthologous genes, single-cell expression experiments, spline fits for pseudo-time analysis and nodes and edges for each of the interaction networks available. Finally, a docker-compose configuration script was written, which contains all relevant configuration parameters in one place to easily deploy the app+sql server services. The App can be accessed at: https://umbibio.math.umb.edu/babesiasc/. The source code for the app is available on GitHub at: https://github.com/umbibio/babesia_dash_app.

## RESULTS

### scRNA-seq analysis reveals the characteristic cyclical pattern of expression in replicating *Babesia* blood-stage parasites

To characterize replication cycle progression in *Babesia spp.*, we performed scRNA-seq on three *Babesia* spp.: *B. bigemina* and *B. bovis* of bovine origin, and *B. divergens* of human origin. To interrogate potential host cell-specific differences, we collected *B. divergens* parasites growing *in vitro* in both bovine and human RBCs. The alignment rates (AR) were over 90% in all species. Downstream processing and alignment detected between 8719 - 12910 cells across the four species. The median number of genes detected per cell varied from 417 to 629. Additional filters were applied using the Seurat R package (51). The number of cells and genes that passed the Seurat cutoffs varied from 3200 - 4000 and 3563 – 3810, respectively across the three species. All alignment metrics are available in **SI** table 1. Expression datasets were combined using the expression of 2548 orthologous genes obtained from reciprocal blast hits and visualized on UMAP and PCA coordinates (**Fig 1**). The projected data reveal a circular pattern shared across all *Babesia* spp. That is characteristic of asexual replication in both *T. gondii* and *Plasmodium spp*. (23, 39, 48). This circular pattern of expression is a consequence of periodicity in gene expression during the replication cycle and the existence of clusters of co-expressed genes. In the UMAP projection, *B. divergens* in human and bovine host cells cluster together, indicating that the effect of different host results in relatively few changes overall versus when compared to the other species.

**Figure 1.**
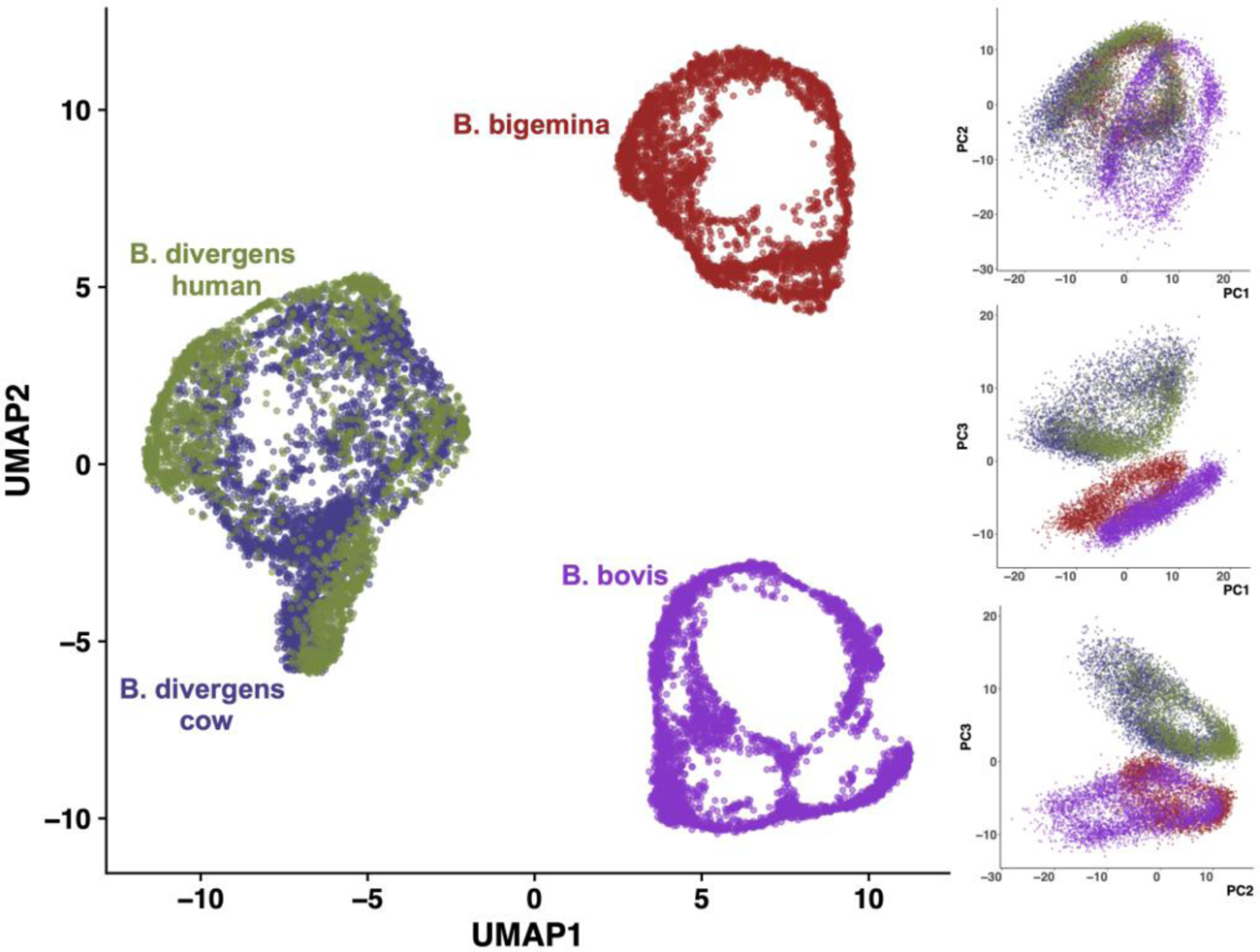
Projection of scRNA-seq data onto UMAP coordinates (Left) and the first three Principal Component (PC) coordinates (Right). B. divergens in human and bovine host cells cluster together.

### Pseudo-time analysis maps the progression of gene expression and mapping transition points in the asexual replicative cycle

To map the progression of gene expression during the replication cycle, we performed a pseudo-time analysis by fitting an elliptic principal curve to the first two PCA coordinates of *B. divergens* (52). Cells were orthogonally projected onto the curve and ordered to mimic pseudo-time (**Fig 2A**). The start point was arbitrarily picked and set to 0 and the end point was set to 12 hr (55, 56), representing the approximate time of a division cycle in *Babesia spp* (37, 56). We partitioned the projected cells along the pseudo-time into 20 minutes time intervals. Cells in each partition were considered as “synchronized replicates”. Expression of genes along the pseudo-time were then used to construct gene expression curves. Genes whose expression did not correlate with the pseudo-time were identified by fitting a Generalized Additive Model (GAM) and calculating the goodness of fit (adjusted p-value < 0.01). Depending on the dataset, a total of 2504 – 2741 genes passed the criteria and were retained for further analysis (**SI** table 1). The excluded genes constitute randomly expressed genes with a nondescript expression pattern indicative of noise.

**Figure 2.**
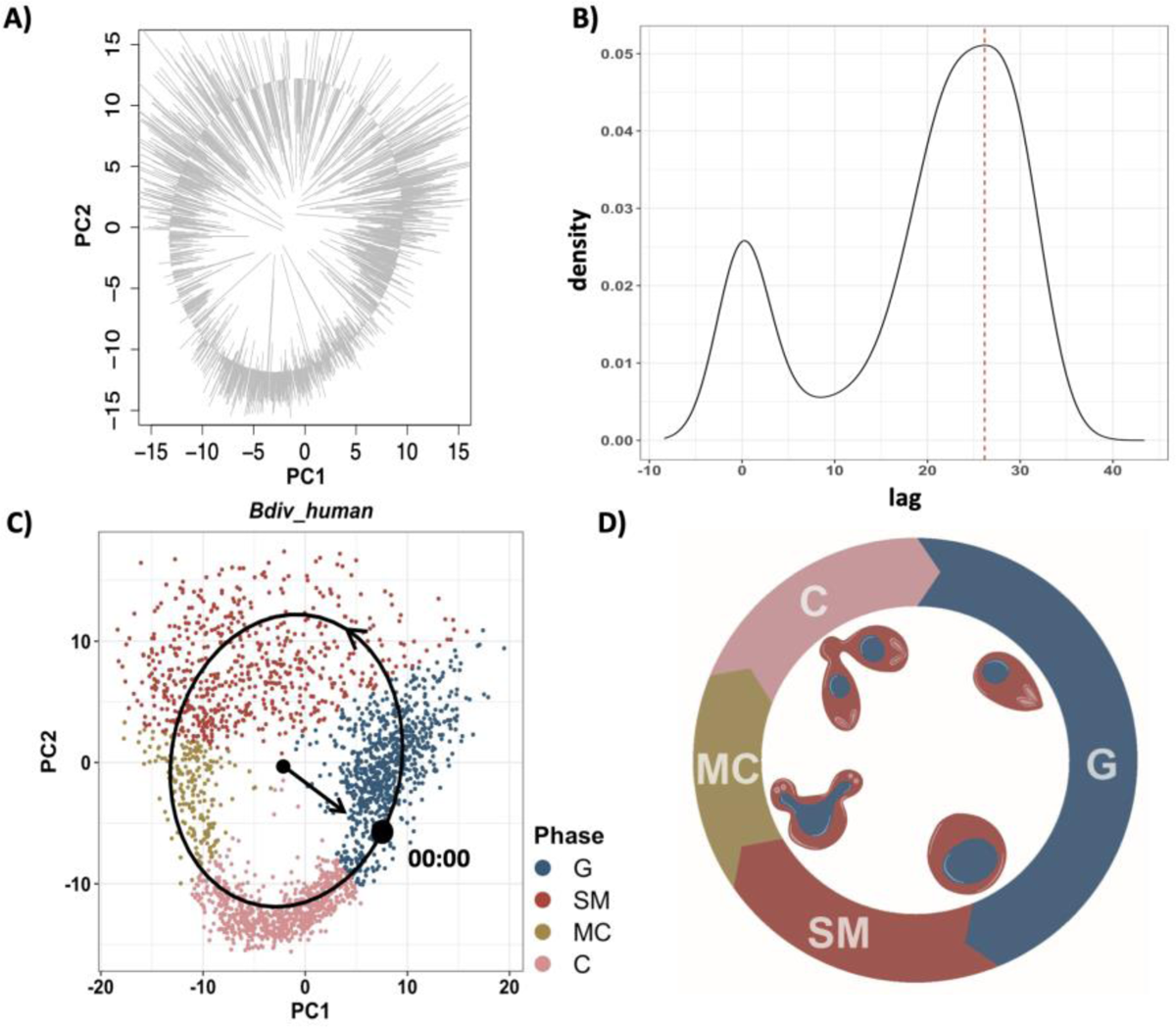
Pseudo-time reconstruction. (A) Cells were orthogonally projected on the elliptic principal curve fitted to the first two PCA coordinates and ordered sequentially using a random start point. Pseudo-time gene curve was constructed by mapping the cell orders to 0-12 h. (B) Distribution of the lag time between pseudo-time gene curves and matched gene curves from synchronized bulk measurements. Lag time was calculated using cross correlation. The bimodal distribution correspond indicates two optimal lag time corresponding to flat curves (lag 0) and peaking curves (indicated red dashed lines). (C) PCA plot of B. divergens in human RBCs. Start time (black dot) was adjusted according to optimal lag time corresponding to the larger peak in panel (B). Black arrow on the principal elliptic curve indicates the flow of time. Transition points were mapped to the PCA coordinates in B. divergens in human RBCs and the colors represent each inferred phase. (D) Schematic representation of the Babesia spp. cell division cycle in RBC.

To adjust the start time and map the direction of replicative cycle progression, we utilized gene expression data from synchronized bulk RNA-seq from *B. divergens* (37). The time course data consisted of 7 bulk RNA-seq measurements of synchronized *B. divergens* parasites over a 12 hr period. The synchronized data consists of 3993 genes, of which 2236 showed significant expression changes over the time (adjusted p-value < 0.01). We calculated the cross correlation of gene expression curves in single cells and the corresponding genes in synchronized bulk experiment and identified the lag-time that maximized the correlation. **Fig 2B** shows the lag time distribution across all genes in *B. divergens*. The two distinct peaks correspond to the optimal shift for constitutively expressed genes (no lag) and peaking genes (lag at 28). The lag corresponding to the second peak was used to adjust the lag-time across all genes and set the start point of the replication cycle clock in single cells (**Fig 2C**).

To identify the transition points through the replication cycle of *Babesia spp.*, we used previously assigned replication cycle markers from *T. gondii*. Although transition points can be inferred computationally, we reasoned that *T. gondii* would be most informative for both its similar binary division modes shared with *Babesia* spp., and because the markers of phase progression for single-cell measurements were assigned using DNA content analysis for calibration (48). Moreover, computationally inferred phases generally agree with *T. gondii* inferred phases (**SI** Fig 1). There is a high correlation between peak expression time of markers and the replicative cycle in *T. gondii* (**SI** Fig 2). The timing of peak expression of the *Babesia* spp. orthologous genes of the top 20 *T. gondii* replicative cycle markers were used to transition time points. There is an overlap between the distribution of peak time expression of S and M as well as M and C makers (**SI** Fig 2). Further information on mapping the transition points can be found in **SI.**

Based on this we opted to define 4 “inferred” phases labeled as: G, SM, MC, and C and marked the transitions to maximize the separation between the phases. These transition time points were used in the pseudo-times to infer developmental phase transition points in *Babesia* spp. **Fig 2C** shows the PCA projection of *B. divergens* in human host cells with the colors indicating the “inferred” phases, accompanied by a schematic depiction of the replication cycle (**Fig 2D**). These results show that by leveraging the geometry of gene expression data in asynchronously dividing parasites, single-cell gene expression can be converted to “pseudo-synchronized” time-course data at fine resolution. This approach can be generally used to overcome some of the limitations of time-consuming and costly time-course experiments.

### Gene expression is highly correlated between synchronized bulk and pseudo-synchronized single-cell

Using the newly inferred phases, we sought to access the correlation of peak time as well as overall similarity of gene curves in the single-cell and synchronized bulk data. First, we utilized a dynamic time warping (dtw) algorithm (57) to align the curves and measured the normalized distance between the curves. **Fig 3A** shows some representative examples of high (left column) and low (right column) alignment for genes that peak (top row) or deplete (bottom row) in inferred SM phase. Next, we calculated peak expression time for each gene in both datasets. The peak assignment was limited to markers of S/M/C phases, where genes tend to have a more defined peak that lasts for a short period of time. The boundary cases of transitioning from C back to G were excluded, as marker genes assigned to G tend to have a flat expression period, followed by depletion at SM phase that slightly upticks in late C and transitions back to G. **Fig 3B** top left shows the correlation between peak expression times for S/M/C genes. The overall normalized alignment distance as measured by the dtw algorithm was categorized into High, Mid, and Low, based on the distribution of the alignment distance (**Fig 3B** top right and bottom panels). Overall, the data shows that there is a high level of agreement between reconstructed pseudo-time in single-cell and the synchronized time-course data, both in peak-time expression as well as overall alignment.

**Figure 3.**
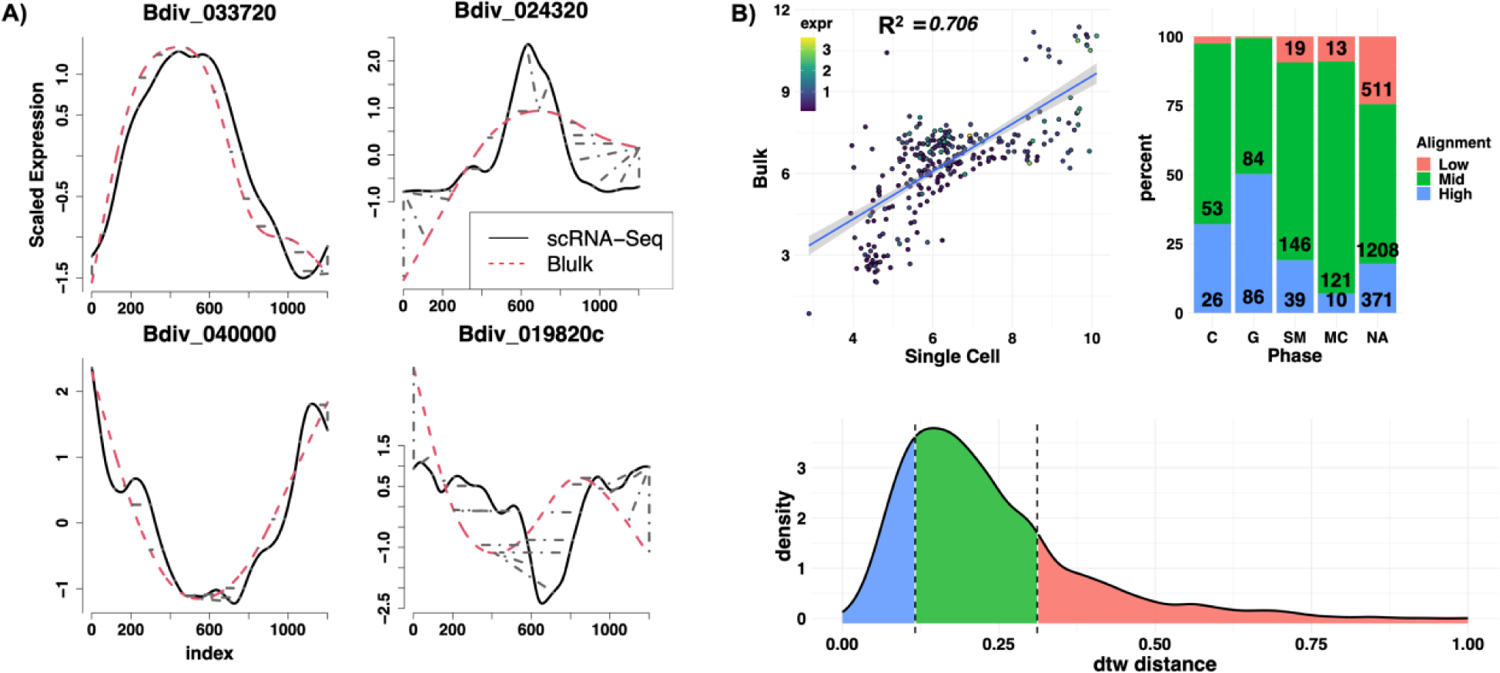
Quantification of alignment of gene curves between synchronized bulk and single-cell. (A) Dynamic Time Warping (DTW) alignment of a sample gene: High alignments (left column), Low alignments (right column). Gray dashed lines indicate the matched time index (warping). (B) Top Left: Correlation between calculated peak times in S/M/C phase, excluding the boundary. Top right: Percent alignment distance categorized into well (distance < Lower 20 percentile) poor (distance > upper 80 percentile) and mid. Curves with no peak contain a higher percentage of poor alignment. Bottom: Distribution of dynamic time warping (dtw) alignment distance between single-cell and bulk. Shaded areas show boundaries alignment categories.

### Analysis of markers of inferred phases

We performed differential gene expressed analysis (DGEA) to identify 1) cross-species conserved replication cycle phase markers, 2) species-specific phase markers (i.e., replication cycle phase markers highly upregulated in one species), and 3) replication cycle phase markers of each species independent of other species (FC > 2 and adjusted p-value < 0.01). The bar plots in **Fig 4A, B, and C** respectively represent the number of conserved replication cycle markers, species specific markers, and independently performed marker analysis in each species. Full lists of gene markers are provided in **SI** table 2. We performed a Gene Ontology Enrichment Analysis (GOEA) to investigate biological functions linked to each set of markers. For this analysis, genes were mapped to their *T. gondii* orthologs and enrichment analysis was performed using available GO terms on ToxoDB. GO term and gene set categories with smaller than 10 genes were excluded from this analysis. Multiple principal significant GO terms are highlighted for the conserved markers in **Fig 4D**. All significant GO term hits can be found in **SI** table 3. The top ranked GO-term for the C inferred phase across the *Babesia* spp. is actin cytoskeleton, including several genes involved in actin polymerization (profilin, Bdiv_003910c; actin depolymerizing factor ADF, Bdiv_021160), as well as myosin A (Bdiv_001770c). These genes are consistent with the cytoskeletal remodeling that occurs during cytokinesis. The SM and MC most highly enriched term relates to the pellicle. In *Babesia* parasites the pellicle is the structural site that organizes budding of daughter cells, thus gene expression would need to occur prior to this process (58). In *T. gondii*, expression of many of the orthologous genes peak during the transition from S to M phase (23), which is consistent with the expression we observe here in *Babesia* spp. A major distinction between SM and MC inferred phase is the distinct upregulation of genes involved in biogenesis and segregation of the Golgi apparatus during SM phase-in mammalian cells this process is known to occur just prior to mitosis (59). This is also in line with previously observed timing of Golgi formation in *T. gondii*, where the Golgi undergoes elongation and segregation prior to the formation of the scaffolding of daughter cells (mitosis) (60). Additionally, genes involved in vesicular transport and vesical formation (clathrin heavy chain, Bdiv_019640c; adaptin N terminal region family protein Bdiv_009600c) were identified in SM phase. Finally, the genes identified as enriched in G phase are generally involved in organelle biogenesis including the apicoplast. In *P. falciparum* apicoplast growth begins in the trophozoite phase prior to DNA replication (61). Further, genes involved in protein folding, including several chaperones and heat shock proteins, are expressed in this stage. Gene involved in DNA replication start expression in late G phase (**SI** Fig 3), suggesting *Babesia spp.* follow a “just-in-time” expression pattern, in this case expressing genes needed during S phase DNA replication during G phase. Further details of phase-based markers can be found in **SI.**

**Figure 4.**
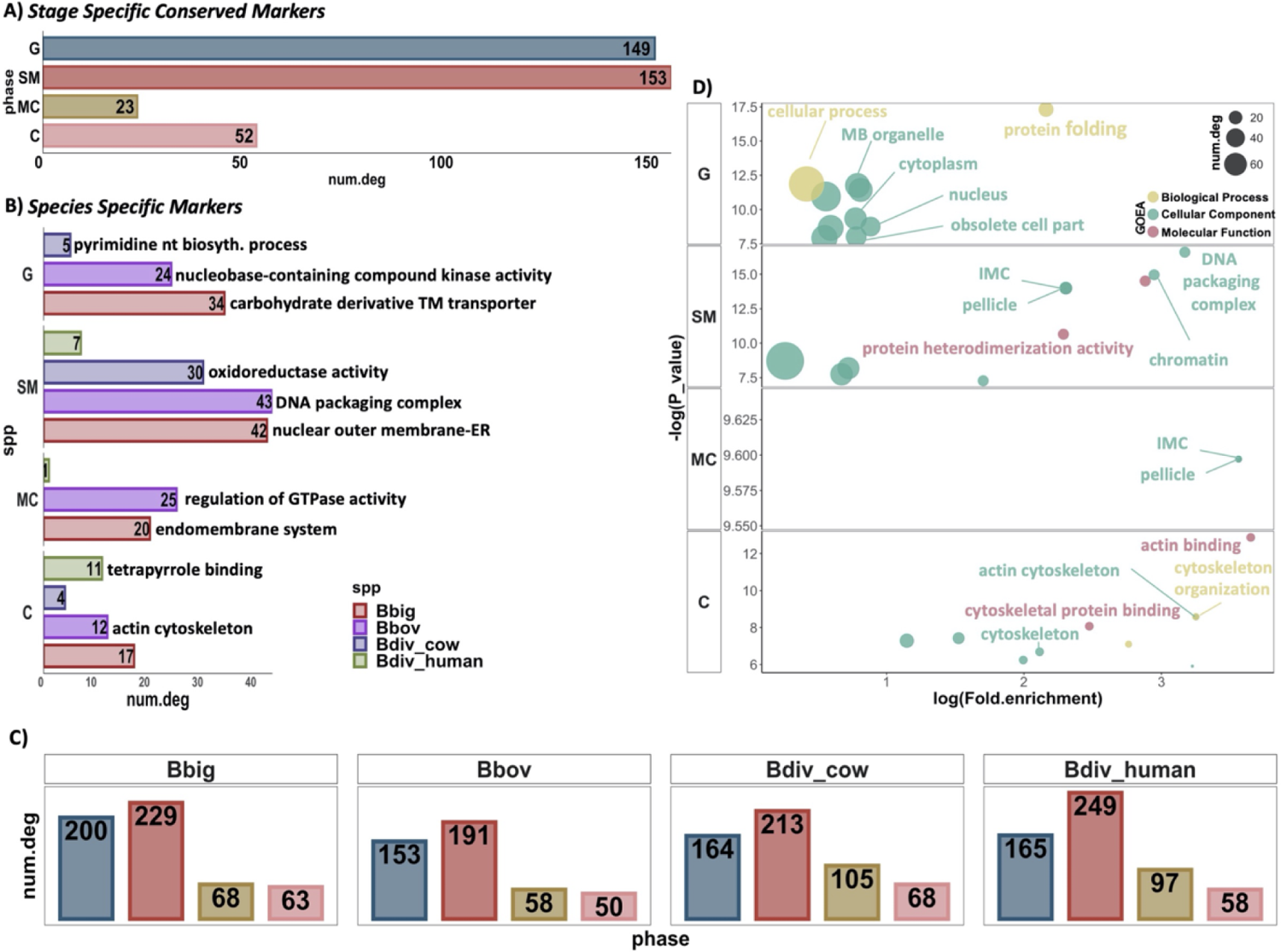
Differential Expression (DE) and Gene Ontology Enrichment Analysis (GOEA). (A) Bar plot shows the total number of conserved markers of inferred replication cycle phases (rows) across all species. (B) Total number of markers of the indicate d phase unique to the indicated species (colors). Most significant (Benjamini < 0.1) GO term associated to the set of markers (each bar) is presented next to the bar. (C) Total number of inferred replication cycle markers in each species independent of other species. (D) Significant GO terms associated with conserved markers of the indicated phase (rows). Colors indicate the GO term category. Note: All markers significant GOEA hits can be found in **SI** tables 2 & 3.

### Analysis of species-specific markers

In addition to conserved markers of replication cycle progression, we also investigated species-specific markers of each inferred state using GO-term enrichment, focusing on the most highly enriched processes (**Fig 4B**). The top most significant GO term for each set of markers is shown next to the bar plot. For each species, a different common thread emerges throughout their respective replicative cycles. Many of the enriched processes involve the endomembrane system and membrane transport in *B. bigemina*. These include nucleotide (ATP) transport (BBBOND_0211990), known to be important in mitochondrial transport in related parasites (62, 63), and several genes with predicted function in the endomembrane system (BBBOND_0309310, BBBOND_0401740, BBBOND_0307820, BBBOND_0102060, BBBOND_0210875, BBBOND_0210880). However, in *B. bovis*, an emphasis on various signaling pathways emerges, including kinase activity (nucleoside diphosphate kinase family protein BBOV_III005290; adenylate kinase BBOV_IV002930) and hydrolase activity (putative GTP-ase activating protein for Arf BBOV_IV012060; GTPase activator protein BBOV_IV007530). For the latter, both enriched genes work to activate GTPases, which are known to be important regulators in other systems (64, 65). Species specific enrichment in *B. divergens* mainly occurs in metabolic pathways, including fatty acid metabolism/pyrimidine biosynthesis (bovine-cytidine diphosphate-diacylglycerol synthase Bdiv_002810c; human-choline/ethanolamine kinase (Bdiv_020970)), oxidoreductase activity (bovine-Bdiv_019910, Bdiv_030660, Bdiv_040430c). Taken together, these results suggest varying levels of dependence on vesicular transport, signaling, and metabolic pathways between *Babesia* spp. Futher description of differences by phase can be found in the SI.

### Progression of gene expression during replication cycle

We sought to cluster and visualize the expression of marker genes during the replication cycle. Markers of each phase were identified in each species independently (**Fig 4C**). We fitted a mixed effect-model with smoothing B-splines to all gene curves and calculated the mean effect along the pseudo-time. **Fig 5** shows the heatmap of scaled expression of mean curves of the inferred replication cycle markers in each species. Genes are ordered by their peak expression time in each phase. Vertical lines show the mapped transition time points, while horizontal lines delineate the markers of each phase. The transition from C back to G consists of genes with “flat” peaks that cross the boundary. To better visualize this, we divided the G phase into Late (G1 L) and Early (G1 E), with G1 E directly proceeding the C phase. The heatmaps show that clusters of marker genes peak at similar times with peak-time expression gradually shifting through the replication cycle. This “just-in-time expression” pattern profile is also seen in other parasite species (23, 24, 66–68), indicating that replication cycle genes are organized into clusters of co-expressed/co-regulated genes that may be functionally related. Heatmap of 377 conserved replication cycle markers across all species (**Fig 4A**) ordered by their progression in *B. divergens* adapted to either human or bovine RBCs show a more similar progression pattern compared to *B. bovis* or *B. bigemina* (**Fig SI 4**), as also seen with the UMAP projection (**Fig 1**).

**Figure 5.**
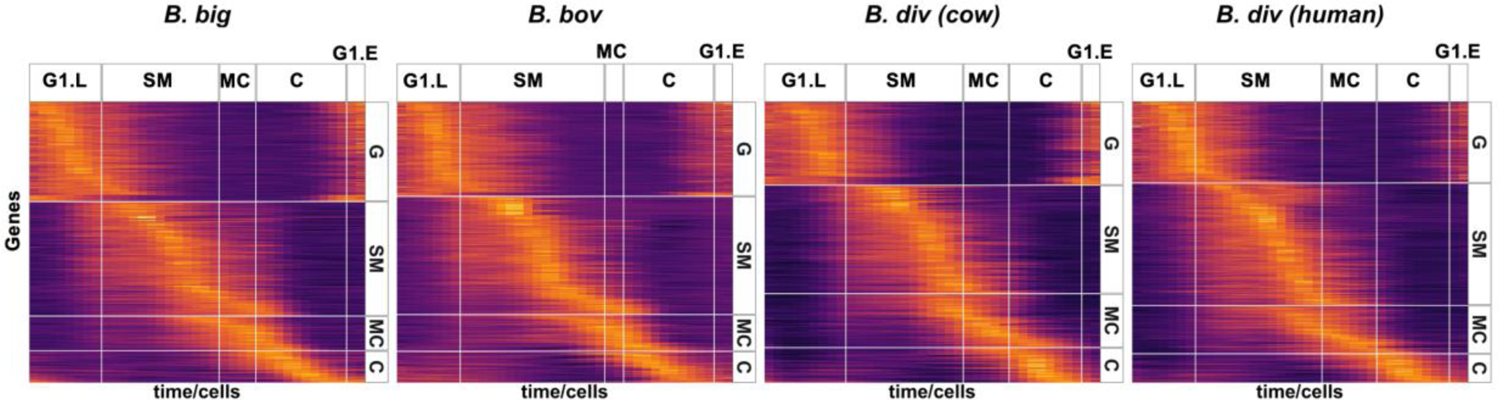
Replication cycle regulated genes transcription profile: Heatmap of mean expression curves of markers of inferred replication cycle phases. Rows represent genes and columns represent pseudo-time/ordered cells. Horizontal facets group the markers of the indicated phase. Vertical facets mark the transition time points. Analysis is done in each species independently (**Fig 4C**). Rows are ordered according to peak expression time.

### Comparative marker analysis of *B. divergens* in human versus bovine red blood cells

To investigate the impact of host cell on the transcriptome, we performed a marker analysis followed by GO term enrichment and identified markers that are uniquely associated to the growth in different host cells. First, we compared the transcriptomes of *B. divergens* in human vs bovine RBCs in each of the inferred phases. This analysis quantifies species-specific host cell impact on the transcriptome. **Fig 6A** top panel shows the expression of the top markers in human (Bdiv_006490c) and in bovine (Bdiv_040280) RBCs. Bdiv_006490c is annotated as an aspartyl protease (*asp3*) that shares homology with plasmepsin X in *P. falciparum*, which is essential for egress and invasion of the parasites (69, 70). Bdiv_040280 is an unannotated protein which contains a mago-binding domain. Mago-domain containing proteins have been shown to be involved in post-transcriptional regulation in other parasites (71). Mago proteins have also been shown to be involved in splicing and trafficking of mRNA (72). The bar plot in **Fig 6A** bottom panel shows the total number of upregulated genes (FC > 1.5 & adjusted p-value < 0.01) specific to *B. divergens* in human RBCs and *B. divergens* in bovine RBCs in each phase. In total, we identified 28 genes across the inferred replication cycle phases that were differentially expressed between *B. divergens* grown in different host RBCs (**SI** table 2). The majority of changes observed occurred in metabolic pathways including lipid metabolism and pyrimidine biosynthesis (refer to SI). These underscore differences in nutrient transport and metabolism based on resident host cell.

**Figure 6.**
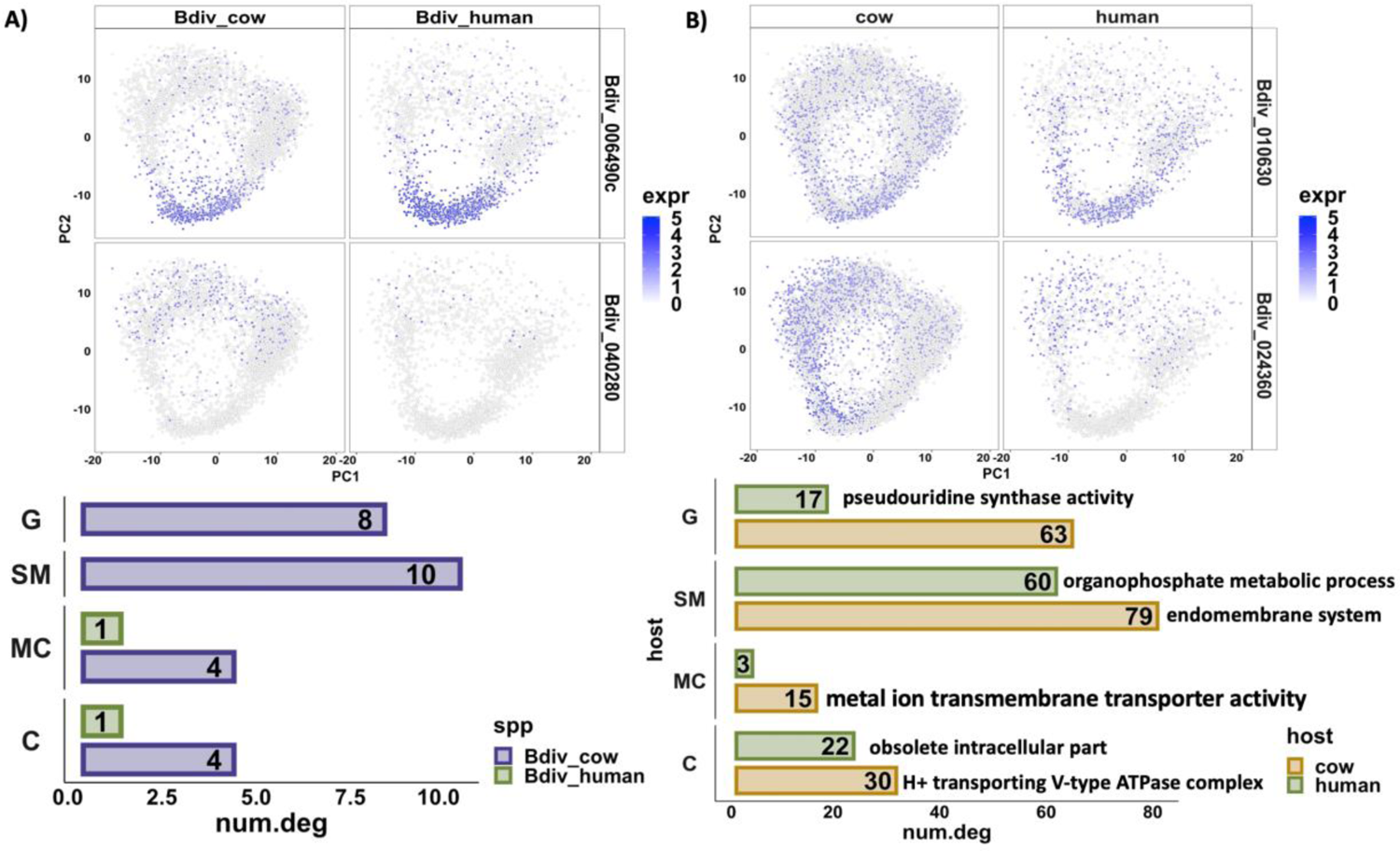
Host specific Differential Expression and GO Term Enrichment Analysis. A) (top), expression of top differentially expressed genes (rows) in B. divergens in bovine RBCs (left) vs B. divergens in human RBCs (right). (bottom) Bar plots show the total number of differentially expressed genes in the indicated phase (rows) in each sample (colors). (B) (top), expression of top differentially expressed genes (rows) in combined B. bigemina, B. bovis, and B. divergens in bovine RBCs (left) vs B. divergens in human RBCs (right). (bottom) Bar plots show the total number of differentially expressed genes in the indicated phase (rows) in each sample (colors). Top most significant GO term associated to each set of markers (Benjamini < 0.1) are presented next to the bar plot.

Second, to quantify the impact of host cell in a species-independent manner we merged all *Babesia spp.* isolated from bovine RBCs and compared the transcriptomes with *B. divergens* in human RBCs for each of the replication cycle phases. **Fig 6B** top panel shows the expression of the top markers in human (Bdiv_010630) and in bovine (Bdiv_024360) RBCs. The human marker Bdiv_010630 is a putative arginyl-tRNA synthetase, while the bovine marker Bdiv_024360 is a conserved hypothetical protein which contains a SNARE-associated Golgi protein domain as well as several transmembrane domains, both of which may be reflective of differences in membrane composition and nutrient availability between the host cells. The bar plot in **Fig 6B** bottom panel shows the total number of upregulated genes in parasites cultured in each host per phase as well as the top significant GO terms. GO terms associated with genes enriched in human adapted *B. divergens* are mostly related to metabolic processes, including nucleotide metabolism (18 genes) and fatty acid metabolism (choline/ethanolamine kinase, Bdiv_020970, Bdiv_020470c), several metabolic pathways including tRNA synthesis (Bdiv_010630, Bdiv_035020c), as well as cytoskeletal dynamics (i.e. profilin, Bdiv_003910c), suggesting possible differences in signaling pathways as well as lipid metabolism in the parasites depending on the resident host cell.

In contrast to the human parasites, in the bovine adapated parasites (*B. bigemina, B. bovis, B. divergens*) the majority of enriched GO terms are related to the Golgi apparatus and endomembrane system and cytoplasmic vesicle formation (Bdiv_005170, Bdiv_009600c, Bdiv_018340, Bdiv_022710), suggesting vesicular transport and protein trafficking may be upregulated in parasites cultured in bovine RBCs. Additionally, there are enriched terms relating to oxidative stress responses (Bdiv_000950) as well as detoxification processes (Bdiv_037500), suggesting the bovine RBCs may be a more oxidatively stressful environment. Taken together these results suggests that there are few differences in cellular processes of parasites propagated in bovine versus human RBCs, which are driven by the host cell environment. Particularly, there appears to be consistent differences in lipid metabolism and vesicular transport. Full list of markers and GO terms are available in SI tables 2 & 3, and are described in **SI**.

### Reconstructing the co-expression network using Gaussian Graphical Models

Presence of clusters of co-expressed genes (**Fig 5**) indicates that shared mechanism may orchestrate the expression of gene clusters and the progression of the replication cycle. To shed light on the interaction between co-expressed genes, we assembled co-expression gene-gene interaction networks using a Gaussian Graphical Model (GGM) (73). For this analysis, we de-trended the time-series expression data by removing the mean trends and utilized a regularized GGM on the de-trended data to construct a gene-gene co-expression network in each species. The sparsity of the networks were controlled using L1 regularization (74) optimized to increase the scale-free property of the network (75–77). Disruption of highly connected hubs in interaction networks causes a major shift in the topology of the network (76). Hence, the hub genes in the network may correspond to genes with essential function in replication cycle progression. We analyzed the overlap of the top hub genes in the networks across all species (**Fig 7A**). There are 10 genes at the intersection of hub genes in all four data sets. Of these 10, three are annotated as surface antigens, 41K blood stage antigen precursor 41-3 (Bdiv_026840c), 12D3 antigen (Bdiv_020800), and 200 kDa antigen p200 (Bdiv_003210), as well as the highly conserved rhoptry associated protein-1 (*rap-1*, Bdiv_025600). Rap-1 is known to be important in host cell invasion, an essential and conserved process in *Babesia* (34, 78–80). **Fig 7B** shows the pseudo time expression of the 10 genes, clustered in two groups. **Fig 7C** shows the inferred interactome of the 10 genes in *B. divergens* (human). Hub nodes in the inferred interactome include *asf-1* (Bdiv_015780c), mitogen activated protein kinase *(mapk:* Bdiv_023270), and calcium-dependent protein kinase 4 *(cdpk4:* Bdiv_024410). ASF-1 has been shown to play an essential role in histone organization and progression through S phase in other systems (81, 82). The MAPK identified shares sequence homology with ERK7 kinase in *T. gondii* (TGGT1_233010), which has a role in stability of the apical complex (83–85). The hub gene *cdpk4* has been previously shown to be essential in egress (37). Interestingly, several hypothetical proteins emerge at the intersection of hub genes, including Bdiv_011410c, Bdiv_024700, and Bdiv_028580c. By determining the interactome of these genes, some inferences about their cellular function can be made.

**Figure 7.**
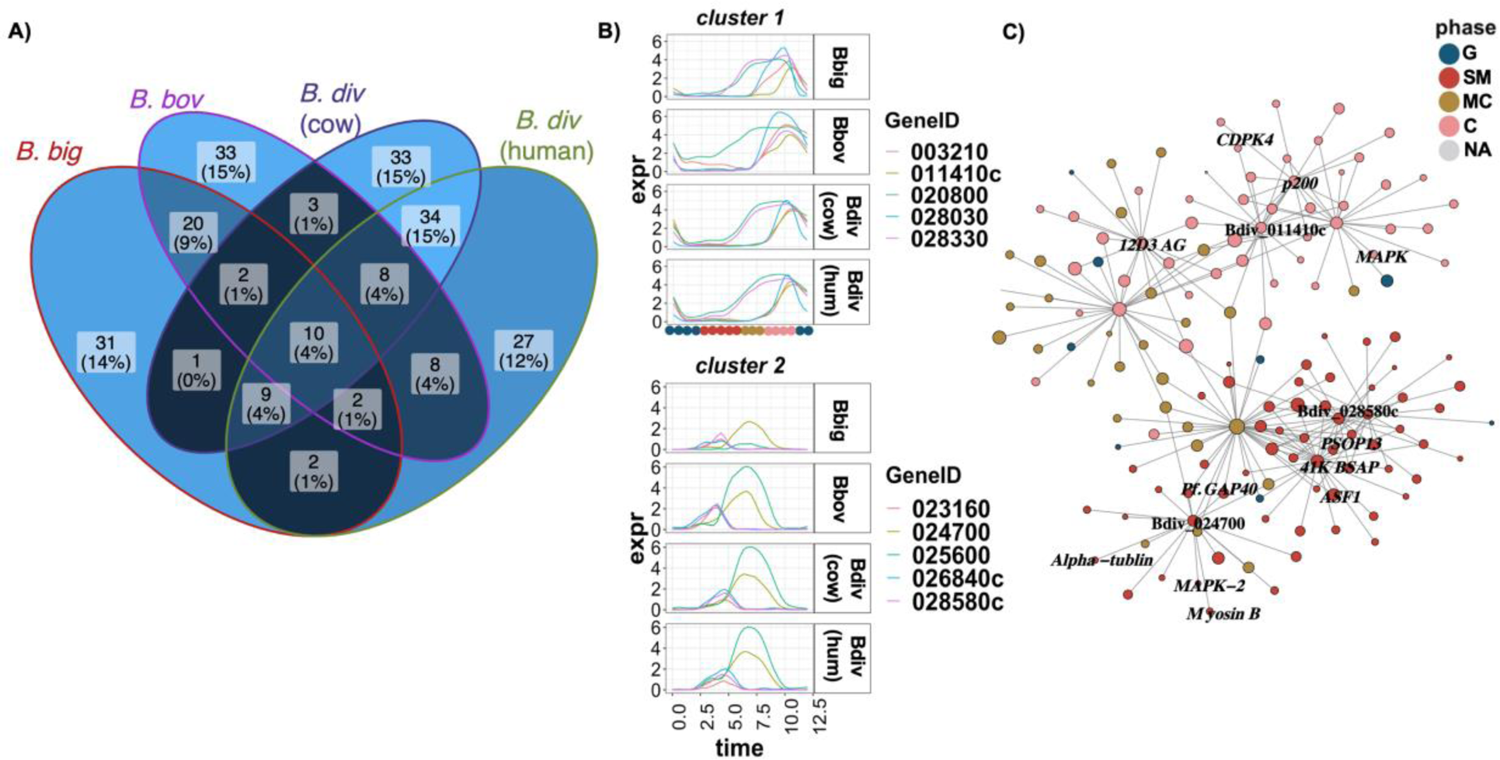
Co-expression Network: (A) Figure shows the number of overlapping genes in the indicated contrast. (B) The expression curves of the 10 conserved hub genes at the intersection grouped into 2 clusters with similar expression profile. (C) The inferred interactome of 10 conserved hub genes in the Babesia divergens (human) co-expression network.

The hypothetical protein Bdiv_028580c shares the majority of its connections with genes in the SM phase (20 of 36 connections). Interestingly, it is connected to an aspartyl protease (*asp6*, Bdiv_022420c) which shares sequence homology with plasmepsin VII of *P. falciparum.* In *P. falciparum*, this aspartyl protease plays an essential role in the invasion of the mosquito midgut in the ookinete stage (86). In addition to this aspartyl protease, this hypothetical protein is also connected to Bdiv_010620, which is orthologous to the secreted ookinete protein, putative (*psop13*) in *B. microti*, and the orthologous gene in *Plasmodium* (PF3D7_0518800) is known to play an important role in transmission (87). Finally, this hypothetical protein is connected to the histone chaperone FACT-L (Bdiv_010540c), which is known to play a key role in male gametocyte development in *P. berghei* (88). FACT-L expression occurs during DNA replication in *P. falciparum* which is consistent with our finding of this hub gene occurring in the inferred SM phase (24, 89). Taken together, these observations suggest a role for Bdiv_028580c in pre-sexual development and suggest this process may be initiated in blood stage *Babesia* parasites. Given the lack of an obvious sexual stage in the presented single-cell data, this suggests parasites may express sexual stage genes in blood stage development to be primed for possible transmission stimuli should they occur. Alternatively, the expression and connection of these markers mainly in the SM inferred phase may indicate an important yet unknown function for these genes in the asexual replicative cycle. Previously, other genes known to be important for *Plasmodium* sporozoite and ookinete development have been found to be expressed in the asexual blood stage of *B. divergens*, for example celTOS (Bdiv_028030) further showing that canonically sexual stage genes are expressed in the replicative cycle (37). Further, genes such *as ama1* have been shown to be important in invasion in multiple *Plasmodium* life cycle stages (90, 91).

The hypothetical protein Bdiv_024700 shares some homology with *ron6* in *T. gondii* (23% identity, TGGT1_297960B), suggesting a possible role in invasion. Bdiv_024700 has several connections to genes important to cytoskeletal arrangement, daughter cell formation, and egress, suggesting a role for the gene in cytokinesis and egress, and most connections occur between SM and MC phase. One such connection is to *mapk-2* (Bdiv_027570c), which is essential in initiation of mitosis and daughter cell budding in *T. gondii* (92). This hub is also connected to the integral cytoskeletal components myosin B (Bdiv_024680) and α-tubulin (Bdiv_038490), as well as Bdiv_020490, which shares sequence homology with Glideosome-associated protein 50 (GAP50, PF3D7_0918000) in *P. falciparum*, which plays a role in organization of the inner membrane complex (93). Indeed, similar cytoskeletal components are required for invasion in *T. gondii* (94). Furthermore, Bdiv_024700 is connected to an aspartyl protease (*asp2*, Bdiv_023140c), similar to plasmepsin IX and X in *P. falciparum*, and has a known role in invasion (37). Together, these data suggest a role in the invasion process for Bdiv_024700.

Finally, looking at the interactome of Bdiv_011410c, most gene connections occur in the inferred C phase. This hypothetical protein is connected to many genes involved in signal transduction, most notably two calcium dependent protein kinases (*cdpk4* Bdiv_024410; protein kinase domain containing protein Bdiv_033990c) and the aspartyl protease *asp3* (Bdiv_006490c), suggesting a possible role in signaling processes which control egress (37).Taken together, co-expression networks tend to connect genes in the same phase and contains few hubs that may have an essential role in the replication cycle.

### Expressional profiles of functionally related gene families

To examine the expressional changes in functionally related genes, we performed a clustering analysis on a list of curated gene families. These include 1) putative transcription factors (TFs), identified by the presence of a DNA binding domain in the sequence, 2) putative AP2 TFs identified by orthology with *P. falciparum*, 3) putative epigenetic factors similarly identified by orthology with *P. falciparum*, and 4) variable erythrocyte surface antigens (VESA) genes (**SI** table 4). The reconstructed expression curves of each gene family in each species were clustered using the dtw algorithm. **Fig 8A** shows the clustering of epigenetic markers with cyclic expression profile in all species. Four clear clusters emerge, with many genes peaking at similar times across all species, although expression pattern of some genes are species specific. Many of the core histone proteins are co-expressed in cluster 1, showing timing of expression occurring mainly during the inferred SM phase. This pattern of coordinated expression of core histone proteins has been observed in related parasites (95). There is also a coordinated expression in the MC phase in cluster 3, including epigenetic factors involved in chromatin organization (Bdiv_023810, Bdiv_024470c, Bdiv_036910c) and histone modification (Bdiv_023060, Bdiv_034310c). Cluster 2 shows peak gene expression of 3 epigenetic genes that occur in C phase, a histone demethylase (Bdiv_012930), histone acetyltransferase (Bdiv_034310c), and a zinc-finger domain containing protein (Bdiv_016880c). Cluster 4 shows broad expression over the replication cycle, and the genes that comprise into this cluster do not fall into similar gene families. **Figure 8B** shows a heatmap representing the presence of each cyclical epigenetic marker in each cluster and species. We next identified the inferred interactome of these markers using the co-expression networks. **Fig 8C** shows that interaction network of epigenetic markers in cluster 1 for *B. divergens.* Interestingly, many of these genes appear as hubs in the network including *asf-1*, which is the top hub gene in *B. divergens* in bovine blood and among the top in *B. divergens* human blood and also appears in the inferred interactome of 10 conserved hubs (**Fig 7C**). ASF-1 shares connections with histone H2A (Bdiv_011310c), histone 2B (Bdiv_011450c), and chromatin assembly factor 1 (*caf-1*) subunit C (Bdiv_013610), each of these occurring in cluster 1 (**Fig 8A**). Identifying this interaction provides support for the validity of the networks generated from these data in *Babesia* spp.

**Figure 8.**
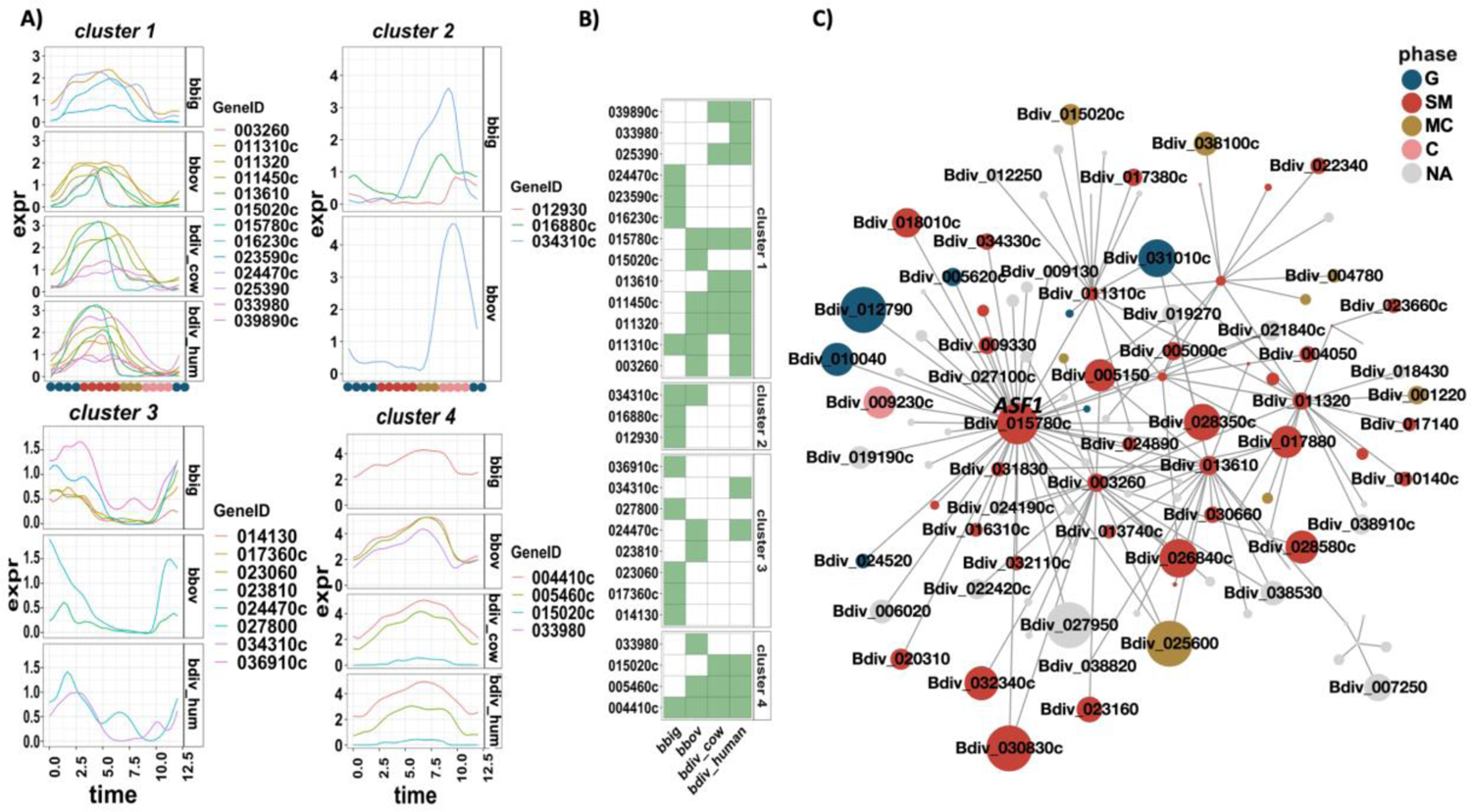
Expression profile of epigenetic markers with <u>a</u> cyclic pattern of expression: (A) The expression curves of epigenetic marker genes clustered into 4 groups according to their expression similarity, <u>split by species</u>. (B) Presence <u>(green)</u> or absence <u>(empty)</u> of the gene (rows) in the indicated sample (column) indicates <u>that the</u> gene is <u>either</u> not cyclically expressed or belongs to a different cluster (<u>033980, 024470c, 015020c, and 034310c).</u> (C) Interaction co-expression network of epigenetic markers genes in B. divergence (human) in cluster 1 (9 total).

The same analysis was performed for TFs with cyclical expression profile, including AP2 domain containing proteins, identified through reciprocal blast with identified transcription factors in *Plasmodium falciparum* (96) (**SI Fig 5**). We identified four clusters of co-expressed genes, which map to the 4 inferred replication cycle phases, with each gene identified by its ID in *B. divergens* (**SI** Fig 5A). Several AP2s domain containing protein appear to show cyclical expression profile in one species. For example, Bdiv_037050 (AP2-G3) peaks at G phase in *B. divergens* and Bdiv_010110c peaks during SM in *B. bigemina.* In contrast, the AP2 domain containing proteins Bdiv_000800c and Bdiv_024900c are expressed at the same time in all species. There appears to be a highly conserved function in the inferred SM phase for Bdiv_024900c-an AP2 domain containing protein with sequence homology to PF3D7_1239200. The expression of this gene is earlier than the expression profile of *P. falciparum*, where it peaks in the later stages of schizogony (97). The most similar ortholog of this gene in *T. gondii* is AP2VIIb-3 (TGGT1_255220), which is implicated in the replication cycle progression into S phase, which is more in line with the observed expression across *Babesia* spp. (98, 99). List of gene cluster IDs are available in **SI** table 5.

### Motif Analysis of the interactome of TFs and AP2

To examine whether the interactome of TF and AP2 family of regulators are transcriptionally co-regulated, we performed a motif search analysis using MEME suite (54) on the promoter genes in the interactome. This analysis was performed on the union of interactome of each AP2 and TF across all species. The analysis identified a significant motif “ACACA” in the promoter of three of the AP2s: Bdiv_015020c, Bdiv_024900c and Bdiv_031830 (**Fig 9A**). These AP2s are orthologous to *P. falciparum* AP2s PF3D7_0604100 (SIP2), PF3D7_1239200 (an unstudied AP2) and Pf3D7_0802100 (AP2-I), respectively. Interestingly, the motif “ACACA” has previously been reported to play a role in DNA replication in *Plasmodium* spp. (100, 101). Moreover, analysis of ATAC-seq regions in *P. falciparum* has identified and associated the same motif with AP2-I Pf3D7_0802100 (102). **Figure 9B** shows the peak time expression of the three AP2s in all species. The AP2s mostly cluster together and seem to peak at S/M phase.

**Figure 9.**
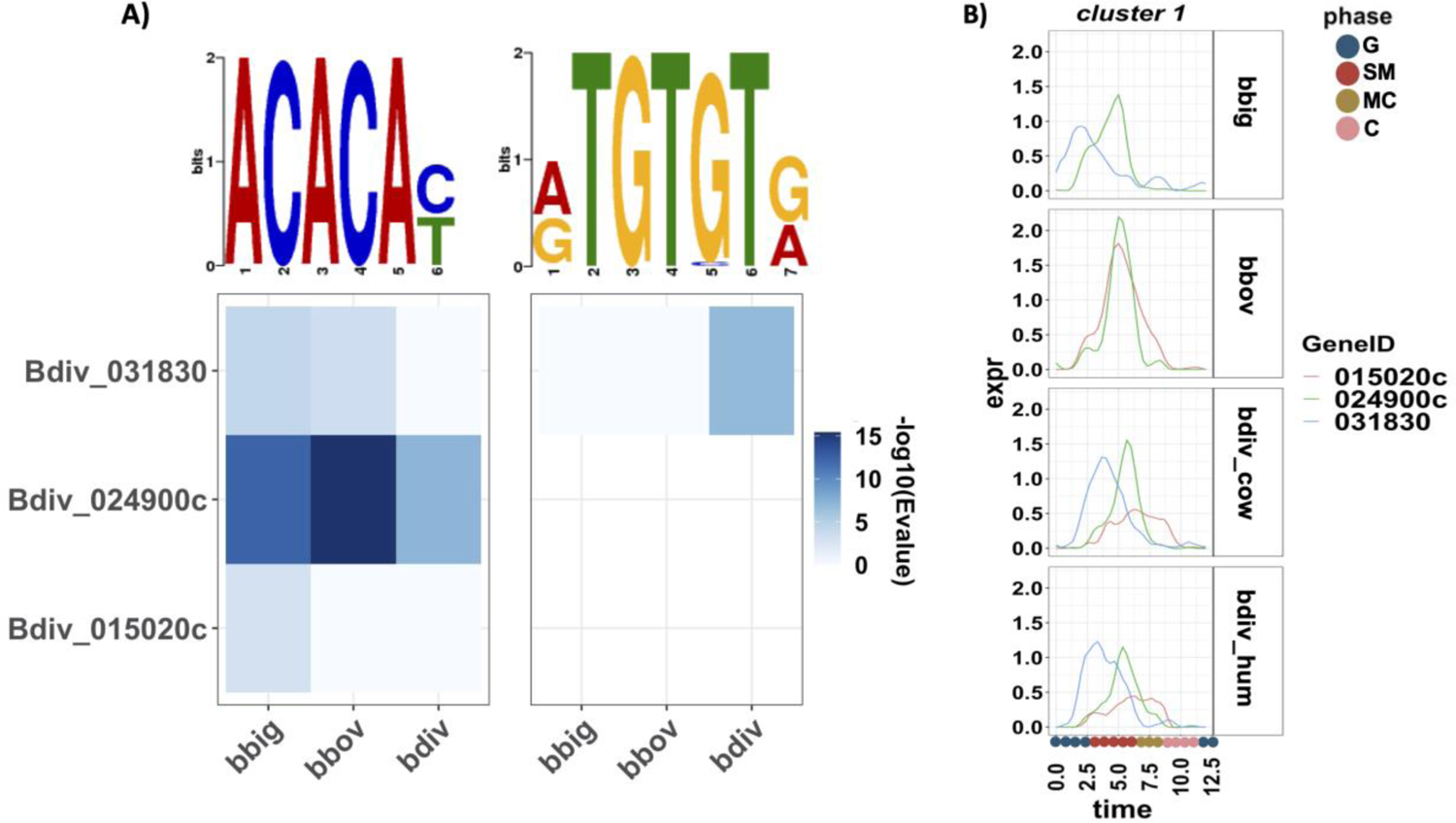
Motif search analysis identified a significant motif in the promoter of three AP2s. (A) The heatmap shows the significance (-log10(Evalue)) of identified motifs with rows corresponding to the gene (AP2) and columns corresponding to the species. (B) Expression curve of the three AP2s.

### Interactive web-app

To facilitate usage, we developed a user-friendly interactive web-app using web dashboard. The App provides functionality to explore and visualize gene expression during the IDC across thousands of asynchronously dividing single cells projected on PCA or UMAP coordinates. Users can examine timing of expression using pseudo-time analysis and perform comparative transcriptomic analysis across the *Babesia* spp. Moreover, users can generate co-expression networks and interactively visualize and explore the inferred interactome of genes. **Fig 10** illustrates the main functions implemented in the App. The App can be accessed at: https://umbibio.math.umb.edu/babesiasc/. The source code for the app is available on GitHub at: https://github.com/umbibio/babesia_dash_app.

**Figure 10.**
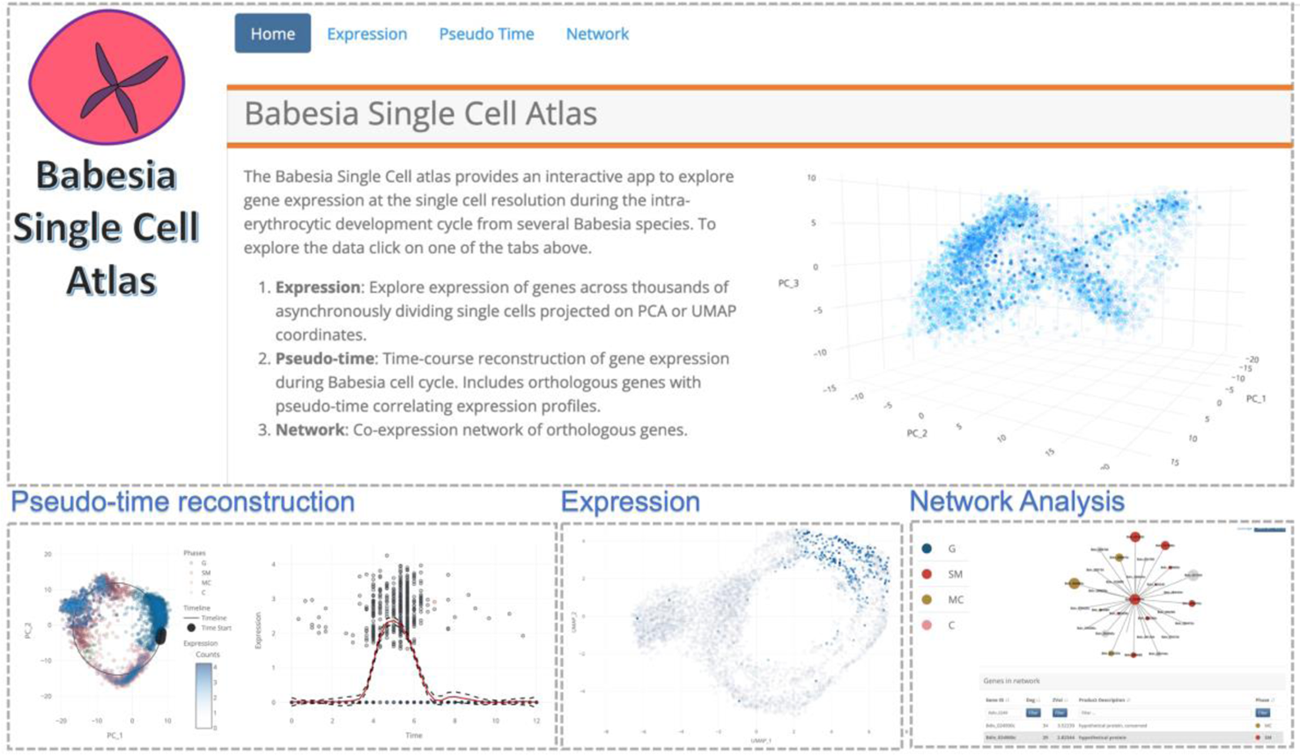
*Babesia* single-cell atlas screenshots: The app provides functionality for pseudo-time analysis, visualizing expression, network analysis, and comparative transcriptomics. URL: https://umbibio.math.umb.edu/babesiasc/

## DISCUSSION

In this work, we present the first single-cell sequencing data in asynchronously replicating *Babesia* parasites and characterized the progression of the replication cycle using newly developed computational approaches. The replication cycle in *Babesia* spp. likely relies heavily on various signaling pathways (reviewed in Elsworth and Duraisingh, 2021). In the asexual replicative cycle of *B. bovis* and *B. bigemina* and many other species, after invasion, the vast majority of parasites will grow, mature, and divide once to form two daughter cells prior to egress; for other species, parasites will divide into four daughter cells (103). Data on the division cycle for *B. bovis* in asynchronous culture suggests that cell division is complete in approximately 5 hr, yet this does not describe the time from invasion to egress (104). However, in a more recent study on synchronized *B. bovis*, this process is approximately 12 hr (56). In contrast, *B. divergens* has a much more complex replicative cycle (55). Indeed, conflicting literature exists suggesting the replication cycle varies from 4-12 hr, however in most cases it appears the minimal time for division is between 4-5 hr (55, 105, 106). Of these, only a single study was conducted on synchronous parasites (55). However, data also shows that the time for the majority of parasites in *B. divergens* to transition from single parasites to paired piriforms is between 10-12 hr (55). Subsequent division cycles are possible in *B. divergens* and the timing of these cycles can vary between 9-14 hr (55). In all studies of the dynamics of *B. divergens* division dynamics, there is a significant range of time for a single division, highlighting the difficulty in generating an exact measurement (55, 105, 106). This variability leads to difficulty in synchronizing *Babesia* parasites, which currently relies on mechanical release of parasites from RBCs using filtration (55, 56), and we have recently shown that parasites may be egress competent at various times in their intraerythrocytic development (37). Unfortunately, no such detailed data on the division time in *B. bigemina* exists, however the replication rate in culture is similar to that in *B. bovis*, and the two parasites follow a similar pattern of dividing from single to double parasites prior to egressing. Based on these data, we opted to set the window of time for the replication cycle at 12 hr for the three parasites (37). Of note, in our analysis we did not distinguish separate clusters of expression for secondary division cycles in *B. divergens*, suggesting the core replication process is conserved and multiple rounds of replication within the same RBC do not require separate gene regulators. Taking advantage of the unique geometry of replicating parasites, we developed a pseudo-time analysis and used the synchronous bulk RNA-seq data in *B. divergens* to calibrate the progression of time in single-cell data. This technique allows us to generate pseudo synchronized time-course data at a fine resolution and reconstruct the expression waves of genes during the replication cycle. Analysis of the data shows high degree of agreement between bulk and single-cell data, demonstrating the ability of single-cell to match (and overcome some of the limitations of) synchronized time-course measurements.

The limitations, challenges, and advantages of scRNA-seq have been extensively reviewed, ranging from comparisons of the most robust tools, understanding drop outs, and discussion of the ability to understand lowly expressed genes (107–114). While scRNA-seq offers extremely fine granularity of cell states, the resolution with which differential gene expression can be detected varies significantly (113). Indeed, in these *Babesia* spp. data sets, we can clearly observe divergence between bulk and single-cell RNA sequencing in the pattern of expression over time in genes that are lowly expressed. For example, protein kinase G (PKG: Bdiv_020500) was recently identified as an essential gene in egress using synchronized bulk RNA-seq and reverse genetics (37), however, this gene is lowly expressed, and the single-cell data has difficulty detecting this gene (**SI** Fig 6). This highlights a key challenge to all methods of differential gene expression: there is often not enough power to confidently characterize lowly expressed genes (113). These limitations are important to consider when attempting to characterize cell populations, especially those which may represent rare cell types.

Using a comparative approach, we utilized data from better studied *T. gondii*, where the replicative cycle phases have been characterized using single-cell and DNA-content (48) to map the replication cycle phases in *Babesia* spp. and identified the markers of “inferred” replication cycle phases. The significant overlap between peak expression time of *T. gondii* based markers indicate that progression of cell division in *Babesia* spp. may not fit well into the canonical model of cell cycle progression (**SI** Fig 2). Mapping transition points in the *Babesia* replication cycle can also be achieved using automatic clustering approaches. We initially used this approach and identified 5-6 phases, which mostly agree with *T. gondii* inferred phases, with little impact on GOEA results (**SI** Fig 1). The progression of gene expression (**Fig 5**) shows a defined transition point between G and S/M phases, and a gradual shift through S/M/C with overlapping transition boundaries. More data, (e.g. time-course DNA content) is needed to map the phases more accurately. However, Gene Ontology indicates the inferred phases of canonical *T. gondii* based phases agree with the known biology of the replication cycle.

The mapping of replication cycle phases allowed us to identify phase regulated marker genes and perform comparative studies between the *Babesia* spp. The analysis revealed conserved and species-specific genes that delineate the inferred states of the replication cycle. Intriguingly, although several of the genes have species specific expression profiles, total number of markers in each phase are comparable and the progression of marker genes during the replication cycle show similar pattern in all species. We also investigated the impact of the host-species on the transcriptome. Of the parasites tested, *B. divergens* is able to grow in both human and bovine RBCs, while *B. bigemina* and *B. bovis* can only be propagated in bovine RBCs. The genes that were identified with differential gene expression in *B. divergens* between bovine and human RBCs indicated changes in expression of genes involved in transcriptional regulation, as well as those potentially involved in invasion and egress. We also identified several differentially regulated aspartyl proteases known to play roles in invasion and egress in various replication cycle phases (37). The differential expression of these genes could be due to differences in host cell receptors, parasite ligands, or host cell membrane composition. A common thread through the differentially expressed genes between human and bovine adapted parasites is the presence of changes in lipid metabolism and vesicular transport. Despite both being mammalian hosts, bovine and human RBCs differ in their composition, size, and deformability (115–118). One striking difference between bovine and human RBCs is the lipid composition of the cell membrane. The composition of bovine RBCs is significantly divergent from most other mammals in that they have low to absent levels of phosphatidylcholine, while having high sphingomyelin levels (116). This stark difference in lipid composition of the host cell membrane may be drivers of the differential expression observed in various genes involved in lipid metabolism; in human-adapted *B. divergens* there are several instances in which lipid metabolism is upregulated in relation to bovine adapted parasites. Previous work has demonstrated that *B. divergens* in only able to synthesize the backbones of phospholipids, and heavily rely on exogenous fatty acids to complete synthesis (119), yet these mechanisms have not been described in parasites cultured in bovine RBCs. Future experiments comparing the differences in these processes between bovine and human adapted parasites are warranted based on the differences observed here.

In contrast, the major unifying difference in bovine parasites is the emphasis on vesicular transport and the endomembrane system. One hypothesis from this observation is that parasites grown in bovine RBCs require increased trafficking to and from the membrane, both to export proteins and scavenge nutrients. The ability of *B. bovis* to cause disease pathology by adhering to the vascular endothelium by altering the RBC membrane surface supports the idea that increased protein export may occur in bovine parasites (120–124). Recently, expression of a multigene family of multi-transmembrane integral membrane proteins (*mtm*) was identified the proteome of the infected RBC membrane, in addition to several other multigene families (*ves1, smorf, tpr-*related), showing this protein is expressed and exported by the parasite (125). These processes in *B. bovis* could account for an increased use of the endomembrane system. However, this cannot fully explain the observed enrichment across the bovine-adapted parasites, as both *B. bigemina* and *B. divergens* do not sequester. However, the case still may be that parasites in bovine RBCs more dramatically alter the host cell. Indeed, when splenectomized calves were infected with stabilites of bovine derived strains of *B. divergens*, an increase in the mean corpuscular volume of the host cell was observed in relation to infection by human derived patient isolates (stabilites), suggesting changes in the membrane architecture. This suggests there is a parasite specific effect on bovine host cells depending on which host (human versus cow) the parasite was originally isolated from (126). Unfortunately, no such studies investigating protein export or RBC architecture exist as of yet in *B. bigemina* (124). However, the evidence from *B. bovis* and *B. divergens* combined with the pattern observed in this study presents an intriguing possibility that protein export is affected by resident host cell.

Our work provides the first comparative single-cell transcriptomic study across three *Babesia* species. To date, comparative genomic studies have focused on gene annotation, understanding variant gene expression, and elucidation of virulence determinants (17, 18, 20, 29, 31–33, 35, 36, 127). In the context of this study, it is worth noting the organization of the genome between *B. bovis* and *B. divergens* share similarity (18). Additionally, *B. bigemina* contains extensive duplications of certain gene families, leading to an increased genome size (32). Genome sequences would suggest similarities exist in general cellular development, and variations arise due to host evasion pathways and differential host tropism. Several comparative studies have sought to understand the expression and structure of variable erythrocyte surface antigens (VESA), both within and across species (32, 36, 127). Consistent with these studies, we also observe a sporadic expression pattern of VESA genes with species specific differences ( **SI** Fig 7).

We also utilized our data set to examine the expression profile of other gene families, including, AP2s, transcription factors and epigenetic markers (96, 128). The analysis revealed distinct clusters of genes with similar peak expression time across species, indicating that shared regulatory mechanisms may be orchestrating the progression of the replication cycle. Of note, we identified that several core histones are co-expressed, which is also observed in related parasites (95). Interestingly, the timing of expression of these core histone proteins differs from that in related parasites: for example histone H3 expression peaks in early schizonts towards the end of DNA replication in *P. falciparum* (25), while in three *Babesia* spp. the expression peaks around 3.75 hr, which is at the beginning of S phase. This difference is likely indicative of the different modes of division on the parasites (schizogony versus binary fission). We were also able to provide strong evidence for the utility of the interactome networks by showing the connection of *asf-1* to histone expression (**Fig 8C**). Interestingly, *asf-*1 appeared throughout our analyses, suggesting an important role of the gene in *Babesia* asexual cycle development. Future studies disrupting *asf*-1 in *Babesia* would provide insight into the nature of the regulation of chromatin formation, as well as reveal any novel functions for the gene in the parasite. These interactomes provide the foundation for future perturbation experiments to understand the directionality of regulatory interactions. Additionally, we were able to identify replication cycle phase dependent patterns of expression of AP2 domain containing proteins (**SI** Fig 5) and were able to identify a conserved motif for a set of these transcription factors (**Fig 9B**).

Finally, to facilitate wider use, we present a web dashboard for interactive exploration of the data. The app provides functionality to examine expression of single genes, comparative analysis of expression profiles and pseudo-time curves across all species, and co-expression network analysis of genes. The app is hosted at: https://umbibio.math.umb.edu/babesiasc/. This resource will allow for novel studies to expand upon and add to analyses of these rich transcriptomic datasets using the interactive interface without the need for expertise in computational methods. The work described here lays the foundation for numerous functional studies to elucidate many facets of *Babesia* biology.

## Data availability

An interactive web-application for visualization and exploration of our datasets can be accessed here: https://umbibio.math.umb.edu/babesiasc/. Source code available on GitHub: https://github.com/umbibio/babesia_dash_app. scRNAseq data (fastq) have been deposited to the Sequence Read Archive (SRA) under the accession number PRJNA803312.

## ACKNOWLEDGEMENT

Thanks to David Allred of the University of Florida for assistance in identifying VESA genes for our analysis and many helpful discussions. Also, thanks to David Degras for helpful discussion regarding network analysis.

