## Supplementary Information for "Comparative single-cell transcriptional atlases of *Babesia* species reveal conserved and species-specific expression profiles"

[†]Joint Authors

**SI Figure 1**

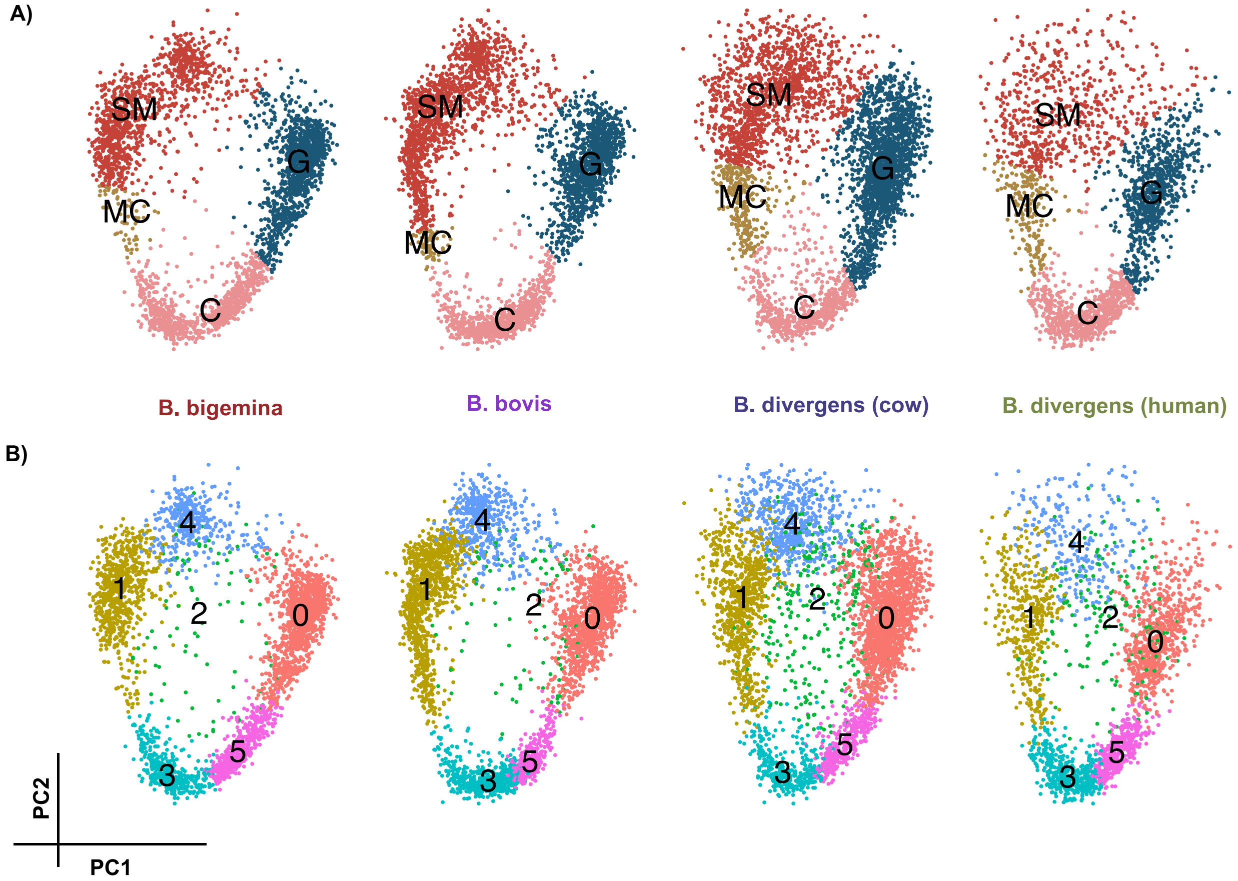


**SI FIG 1.** Figure shows the expression data in all species projected on PCA coordinates. Colors indicate (A) inferred cell cycle phases using *T. gondii* markers, and (B) cell clusters automatically identified using graph-based clustering. Total number of clusters was set to 6.

**Extended information on mapping transition points in the asexual replicative cycle**

**SI Fig 2** shows the distribution of the peak expression time of the *Babesia spp.* orthologous genes of the top 20 *T. gondii* replicative cycle markers, scaled to 0-12 h. The distribution of markers of the G phase is distinct with low overlap with other phases, whereas markers of S phase and early M phase, as well as M phase and early C phase overlap significantly. As such transition points in canonical S and M as well as M and C phases are not readily discernible by timing of expression alone.

### SI Figure 2

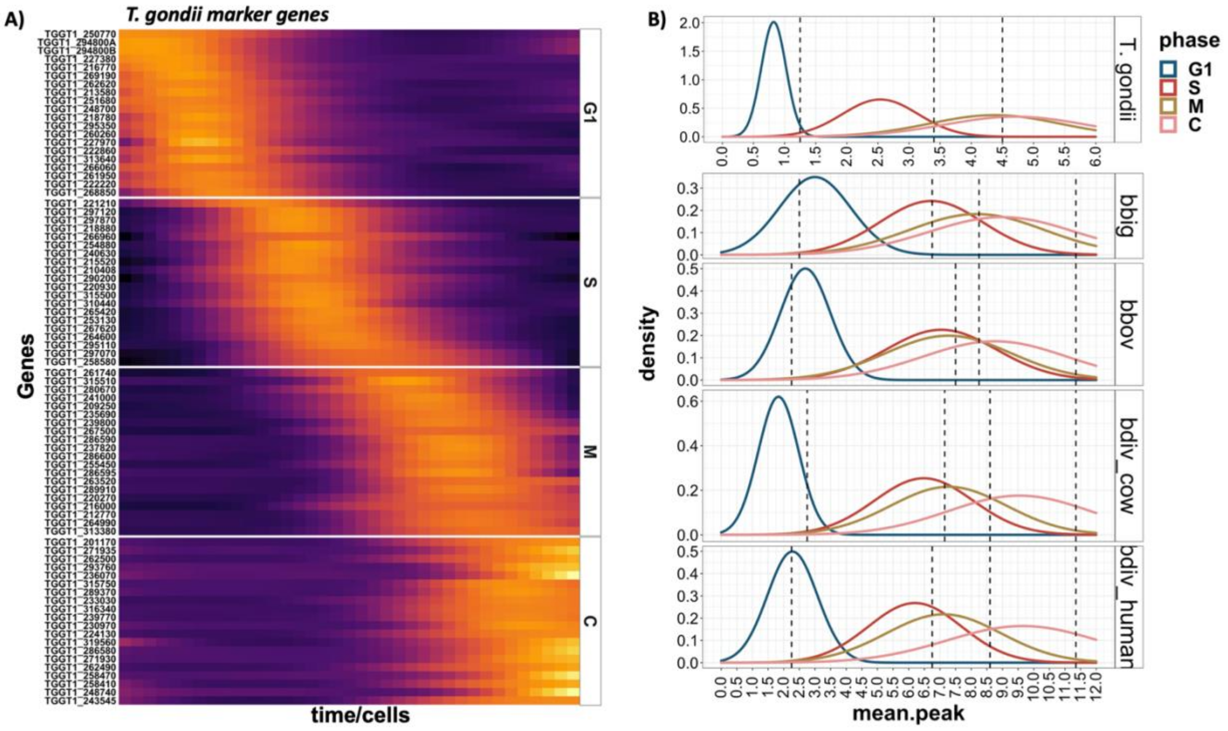


**SI FIG 2.** (A) Heatmap of mean expression curves of top 20 cell cycle phase marker genes in *T. gondii*. Genes with similar expression curve are clustered together. Genes are split according to their identified cell cycle phase from Xue et.al. Work. (B) Distribution of the peak time expression of top 20 *T. gondii* cell cycle markers in *T. gondii* (top panel) and all Babesia species. The peak time curves are colored based on indicated cell cycle phase. Vertical dash lines represent the transition time points.

### SI Figure 3

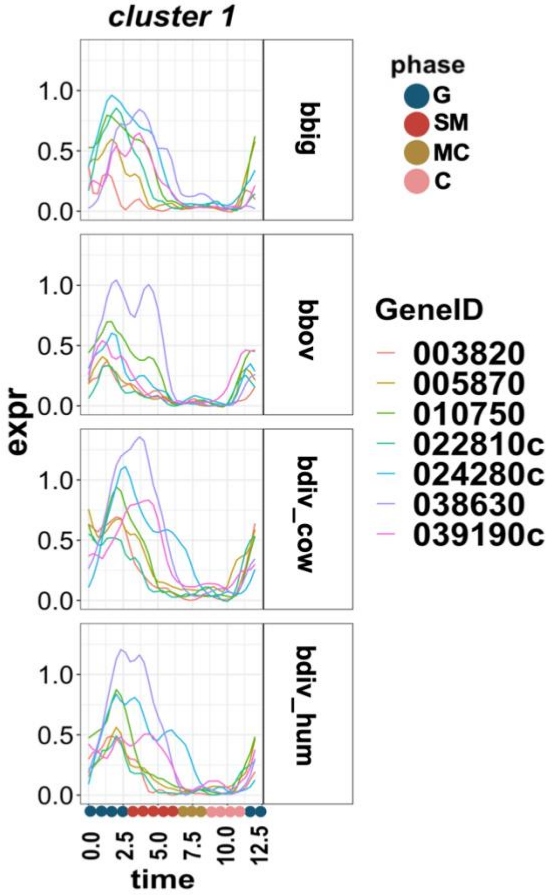


**SI FIG 3.** Expression profile of genes involved in DNA replication that are markers of inferred G phase, split by species.

**Extended analysis of species-specific markers**

In addition to conserved markers of replication cycle progression, we also investigated species-specific markers of each inferred state using GO-term enrichment, focusing on the most highly enriched processes. In the inferred G phase of the three parasite species, the main differences occurred in metabolic processes and nutrient scavenging. In *B. bigemina,* there was strong enrichment for various transmembrane transport processes, the most highly ranked being nucleotide (ATP) transport (BBBOND_0211990), known to be important in mitochondrial transport in related parasites (1, 2). In *B. bovis*, the top enriched processes were more variable than in *B. bigemina*, including actin dynamics (profilin, BBOV_II006000) and kinase activity (nucleoside diphosphate kinase family protein BBOV_III005290; adenylate kinase BBOV_IV002930). In *B. divergens* propagated in bovine erythrocytes the most enriched divergent processes were related to fatty acid metabolism/pyrimidine biosynthesis (cytidine diphosphate-diacylglycerol synthase Bdiv_002810c) and iron transport (GTP-binding protein engA Bdiv_017520c). Taken together, each of these suggest there are subtle differences in metabolism between the parasites in the inferred G phase. There were no specifically enriched terms in *B. divergens* cultured in human erythrocytes.

In the inferred SM phase, the most highly enriched GO-term for *B. bigemina* is nuclear outer membrane-endoplasmic reticulum membrane network (BBBOND_0309310, BBBOND_0401740, BBBOND_0307820, BBBOND_0102060, BBBOND_0210875, BBBOND_0210880), suggesting a species-specific increased role for the endomembrane network during this phase. In *B. bovis,* protein-DNA complex is enriched (BBOV_III007560); these genes are both histone proteins, suggesting an increased importance of chromatin structuring during SM phase for *B. bovis.* Finally, the most highly enriched processes for *B. divergens* is oxidoreductase activity (bovine RBCs) (Bdiv_019910, Bdiv_030660, Bdiv_040430c). The observed differences in *B. divergens* are likely driven by the host cell: for bovine RBCs the enriched genes suggest an increased role of the oxidative pentose-phosphate pathway, while in human RBCs a shift towards expression of proteins needed for post-transcriptional activity. While parasites clearly all undergo DNA replication during the inferred SM phase, these differences suggest that key processes differ during this phase.

In the inferred MC phase there again appears to be an enrichment for activities involved in the endomembrane system in *B. bigemina*. In *B. bovis* there is an enrichment for regulation of hydrolase activity (putative GTP-ase activating protein for Arf BBOV_IV012060; GTPase activator protein BBOV_IV007530) - both enriched genes work to activate GTPases, which are known to be important regulators of mitosis in other systems (3, 4). The orthologs of these genes in *T.*

*gondii* have recently been shown to localize to the nucleus, with the latter of the two appearing to specifically act in the nucleolus, and may play a role in chromatin formation (5). Interestingly for *B. divergens* there are very few specific markers for MC phase. Indeed for *B. divergens* (bovine) there were no enriched terms, and only a single gene for human- choline/ethanolamine kinase (Bdiv_020970)- suggesting that there is little deviation in the MC phase outside of the conserved process in *B. divergens.*

Finally, for the inferred C phase, there are no significantly enriched species-specific markers for *B. bigemina.* However, for *B. bovis* there is a marked enrichment for proton-transport (proteolipid subunit c BBOV_II002740; vacuolar ATP synthase subunit d BBOV_II001540). In *B. divergens* (bovine) a single gene is enriched – protein disulfide-isomerase (Bdiv_013520c), which in other systems resides in the endoplasmic reticulum and performs an essential function in forming disulfide bonds during protein folding (6). In human adapted *B. divergens*, oxidoreductase activity appears to be enriched, driven by upregulation of Bdiv_020470c (cytochrome b5-like Heme/Steroid binding domain containing protein). Together, these results suggest there are only subtle differences in parasites during cytokinesis, which is likely a highly conserved process (7).

### SI Figure 4

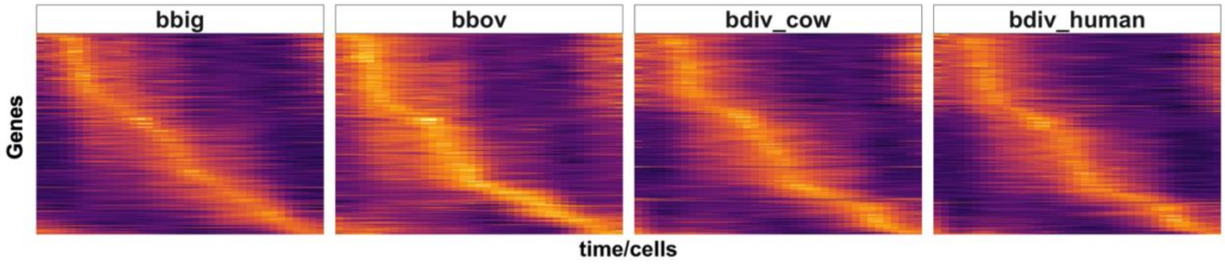


**SI FIG 4.** Heatmap of gene expression curves of 377 conserved replication cycle markers across all species (**Fig 4A**). Rows are ordered according to peak expression time in *B. divergence* in human RBCs.

### Extended Comparative marker analysis of *B. divergens* in human versus bovine red blood cells

In total, we identified 28 genes across the inferred cell cycle phases that were differentially expressed between B. divergens grown in different host RBCs. Of those, eight were from G inferred state, ten in SM, five in MC, and five in C. In nearly all inferred states, the major upregulation of genes was in parasites grown in bovine RBCs. In G, two upregulated genes were involved in ubiquitin related processes (Bdiv_039000c, Bdiv_017290). Additionally, several genes upregulated in bovine versus human propagated B. divergens in inferred G state were related to transcription and DNA replication (Bdiv_005180c, Bdiv_019130, Bdiv_030460). Progressing into the SM inferred state, again all upregulated genes were observed in parasites cultured in bovine RBCs. These genes were involved in metabolic processes including lipid metabolism (3-oxo-5-alpha-steroid 4-dehydrogenase family proteins; Bdiv_039610c), pyrimidine biosynthesis (orotidine 5'-phosphate decarboxylase, Bdiv_024970c) and membrane transport (formate/nitrite transporter family protein, Bdiv_007750c). Additionally, there was enrichment for genes related to membrane components (Bdiv_013030c, Bdiv_007750c, Bdiv_017140). The ortholog of the hypothetical protein Bdiv_013030c localized to the Golgi in T. gondii, while the ortholog of Bdiv_007750c appears to be in the plasma membrane (5). These again seem to underscore differences in nutrient transport and metabolism based on resident host cell. In the MC inferred state, of the five differentially expressed genes, one was specific to human adapted B. divergens - a putative phosphotransferase (Bdiv_020970). The rest of the upregulated genes occurred in bovine adapted parasites, including those involved in cytoskeletal arrangement (Bdiv_038490, Bdiv_016060), and interestingly histone H2Bv (Bdiv_005460c).

**SI FIG 5**

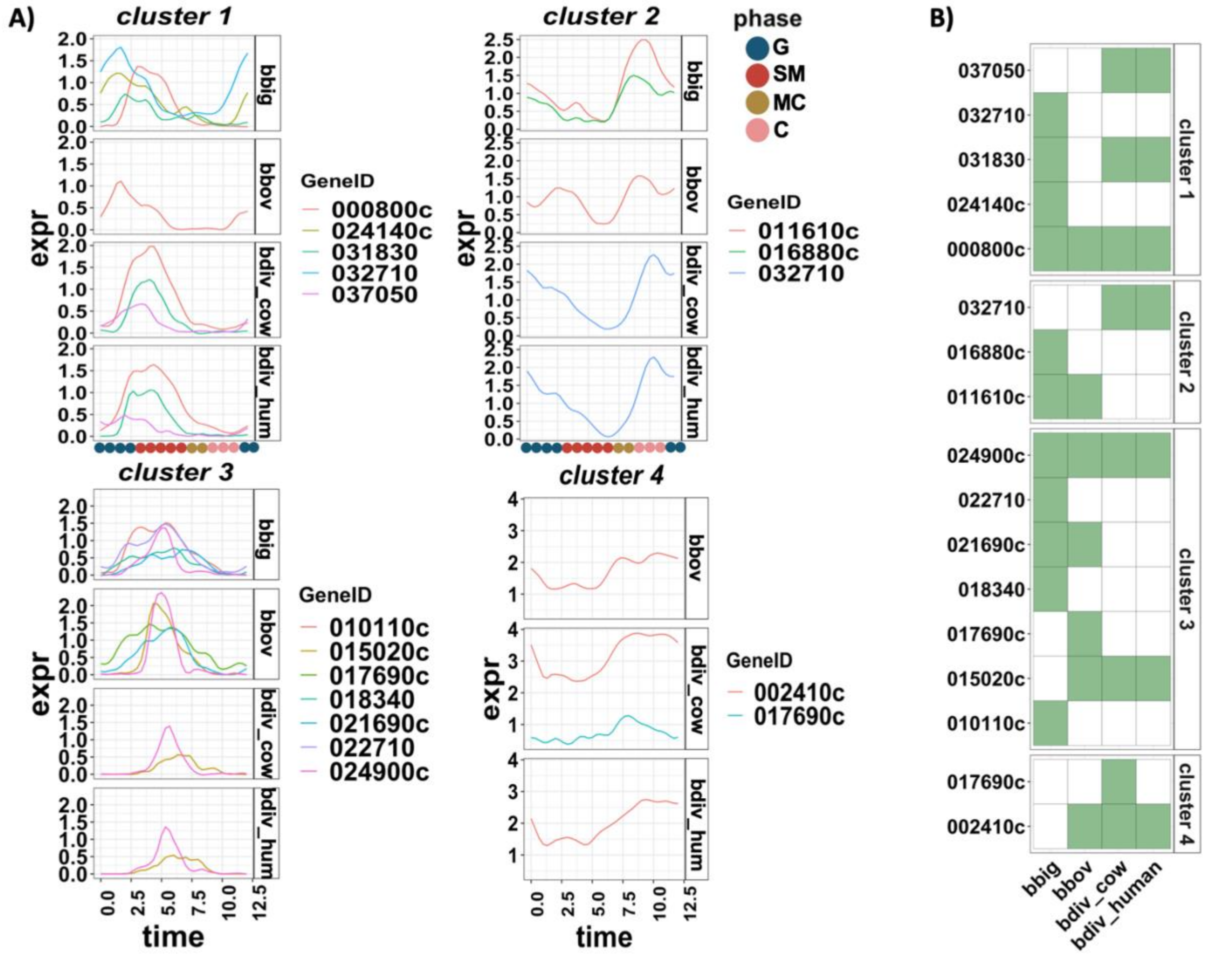


**SI FIG 5.** Expression profile of Transcription Factors with cyclic pattern (TFs): (A) The expression curves of TFs clustered into 4 groups according to their expression similarity, split by species. (B) Presence or absence of the gene (rows) in the indicated sample (column). In the heatmap, white indicates that gene is not cyclically expressed in the species, while green indicates that the gene is cyclically expressed in the related species. There are two genes (032710 and 017690c) that switch the clusters. Note: This is clusters of TFs with cyclic expression profile, not all TFs.

**SI Figure 6**

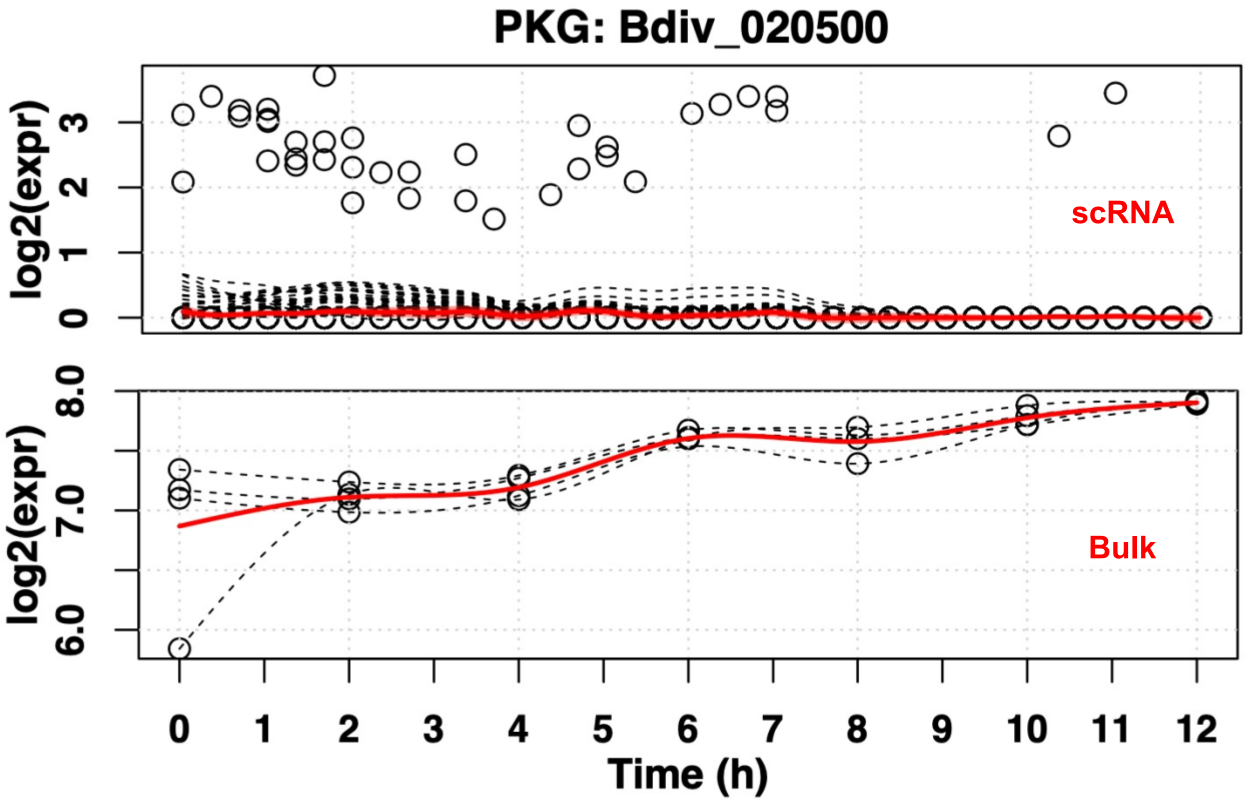


**SI FIG 6.** Expression of PKG in single cell (top) and synchronized bulk (bottom)

### SI Figure 7

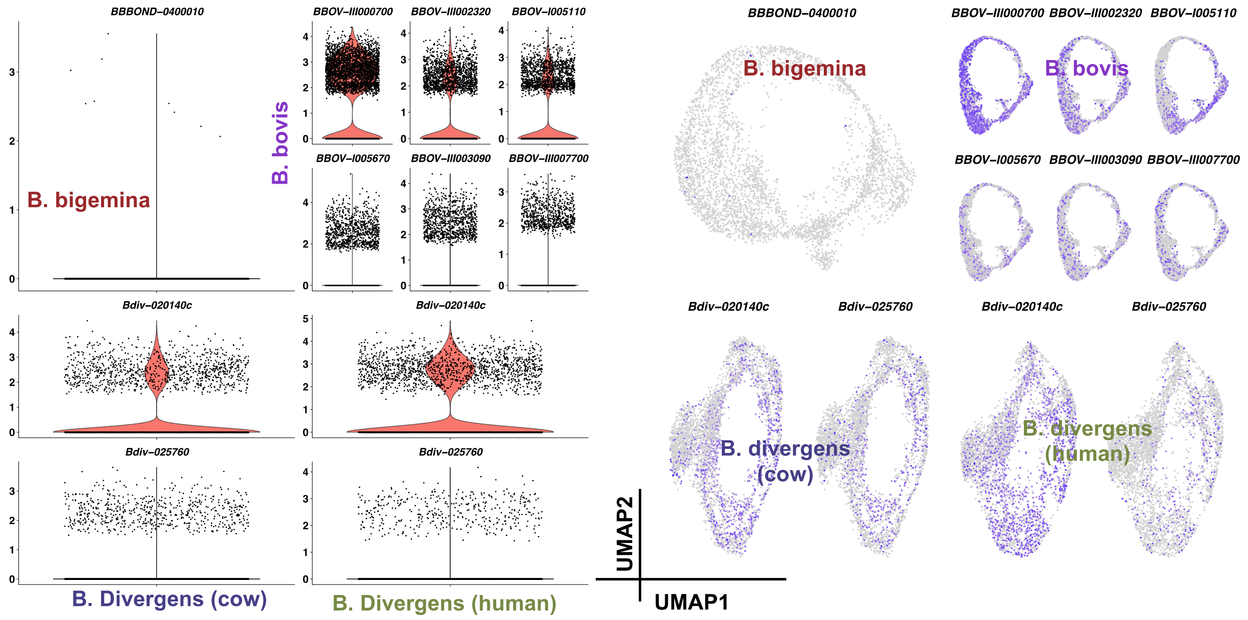


**SI FIG 7.** Expression of highly expressed (95 percentile) VESA genes: (left) violin plots, (right) UMAP projection.
